# ATP dependent polymerization dynamics of bacterial actin proteins involved in *Spiroplasma* swimming

**DOI:** 10.1101/2021.04.07.438887

**Authors:** Daichi Takahashi, Ikuko Fujiwara, Yuya Sasajima, Akihiro Narita, Katsumi Imada, Makoto Miyata

## Abstract

MreB is a bacterial protein belonging to the actin superfamily. It polymerises into an antiparallel double-stranded filament that generally functions in cell shape determination by maintaining cell wall synthesis. *Spiroplasma eriocheiris*, a helical wall-less bacterium, has five classes of MreB homologs (SpeMreB1-5) that are likely to be involved in swimming motility. Here, we investigated the structure, ATPase activity, and polymerisation dynamics of SpeMreB3 and SpeMreB5. SpeMreB3 polymerised into an antiparallel double-stranded filament, and SpeMreB5 formed sheets, including the antiparallel filament, upon the binding of a nucleotide. SpeMreB3 showed slow P_i_ release owing to the lack of an amino acid motif conserved in the catalytic centre of MreB family proteins. Our crystal structures of SpeMreB3 and analyses of the mutant variants showed that the amino acid motif most likely plays a role in eliminating the proton of the nucleophilic water for ATP hydrolysis. Our sedimentation assay suggests that SpeMreB3 has a lower polymerisation activity than SpeMreB5, while their polymerisation dynamics are qualitatively similar to those of other actin superfamily proteins, in which ATP hydrolysis stabilises the filament, and P_i_ release leads to depolymerisation.

## Introduction

MreB is an actin superfamily protein found in elongated walled bacteria (1, 2). Previous studies have reported that MreB polymerises into an antiparallel double-stranded filament depending on the binding of nucleotides, such as ATP and GTP (3-7). The conformational change of MreB upon polymerisation induces the rearrangement of the conserved glutamate and threonine residues interacting with the putative nucleophilic water for γ-P_i_ of the bound nucleotide, thereby facilitating nucleotide hydrolysis (4, 8). Although the residues important for nucleotide hydrolysis are conserved in many MreBs (4, 9), the hydrolysing mechanism and the role of hydrolysis in polymerisation dynamics remain unclear.

MreB functions as a scaffold of an “elongasome “ complex for the synthesis of the peptidoglycan layer, the bacterial cell wall during the cell elongation phase. Through the MreB function, the bacterial cell is arranged in a rod shape (2, 10). MreB forms a filament on the cell membrane with very slow subunit turnover. The filaments move in a direction perpendicular to the cell axis, coupled with peptidoglycan synthesis rather than polymerisation dynamics (11, 12). However, some MreBs play distinct roles besides cell wall synthesis, such as *Myxococcus xanthus* MreB, which drives cell gliding (13) and *Helicobacter pylori* MreB, which is involved in chromosome segregation and urease activity (14). Recently, new MreB proteins involved in swimming motility have been identified in *Spiroplasma* species (9, 15).

*Spiroplasma* belongs to the class Mollicutes, which evolved from the phylum Firmicutes, including *Bacillus subtilis* (16-18). They have a helical-shaped cell lacking the peptidoglycan layer and show unique swimming motility in which the cell moves forward by transmitting helicity switching along the cell axis to rotate the cell body. This motility is unrelated to major bacterial motilities, such as flagellar and pili motilities (16, 19-22). Helicity switching and its transmission are likely to be caused by conformational changes in the internal helical ribbon structure along the entire cell axis (19, 23-25). The ribbon structure is thought to be composed of a fibril, a cytoskeletal protein unique to *Spiroplasma* (19, 23, 24), and five classes of MreB proteins (MreB1-5) (25-27). A recent structural study of *Spiroplasma citri* MreB5 (SciMreB5) showed that SciMreB5 has a canonical actin fold and forms filaments (3).

Based on the sequence similarity among the five MreBs, they were divided into three functional groups: MreB1&4, MreB2&5, and MreB3 (Fig. S1A) (26). A recent study proposed the functions of each MreB group by a heterologous expression system, as follows: MreB1 and/or MreB4 form a static backbone interacting with fibril filaments along the cell; MreB2 and/or MreB5 actively polymerise and depolymerise to change the conformation of the ribbon structure; and MreB3 anchors MreB1 and/or MreB4 onto the cell membrane via its amphipathic helix (Fig. S1B) (26). MreB5 is essential for helical cell shape and swimming motility in *S. citri* (3). To understand the swimming mechanism, it is necessary to clarify the molecular features of these MreBs, including their structure, ATPase activity, and polymerisation dynamics. Here, we studied MreB3 and MreB5 in *Spiroplasma eriocheiris* (SpeMreB3 and SpeMreB5), a model organism for studying *Spiroplasma* swimming (19, 20, 23, 28, 29). SpeMreB3 polymerises into an antiparallel double-stranded filament, and SpeMreB5 forms a sheet which contains the antiparallel filament upon nucleotide binding. Our crystal structures of SpeMreB3 and P_i_ release measurements suggest that SpeMreB3 lacks an amino acid motif for ATP hydrolysis conserved over the MreB family proteins, resulting in low ATPase activity. These differences between SpeMreB3 and SpeMreB5 are likely essential for their distinct cellular roles. Our data also suggest the following two molecular features of MreBs: an ATP hydrolysis mechanism of MreB family proteins, including a possible proton transfer pathway, and the polymerisation dynamics of SpeMreB3 and SpeMreB5, which are qualitatively similar to other actin superfamily proteins (30)

## Results

### Nucleotide binding induces polymerizations of SpeMreB3 and SpeMreB5

Purified monomeric SpeMreB3 and SpeMreB5 (Fig. S2) were incubated with or without Mg-ATP for 3 h and observed by negative-staining electron microscopy (EM). In the presence of Mg-ATP, SpeMreB3 polymerised into double-stranded filaments without helicity (Fig. 1A). In contrast, SpeMreB5 formed sheet structures composed of multiple protofilaments (Fig. 1B). However, filamentous structures were not observed in the absence of Mg-ATP (Fig. S3A and B), indicating the nucleotide dependence of the polymerisation.

**Figure 1.**
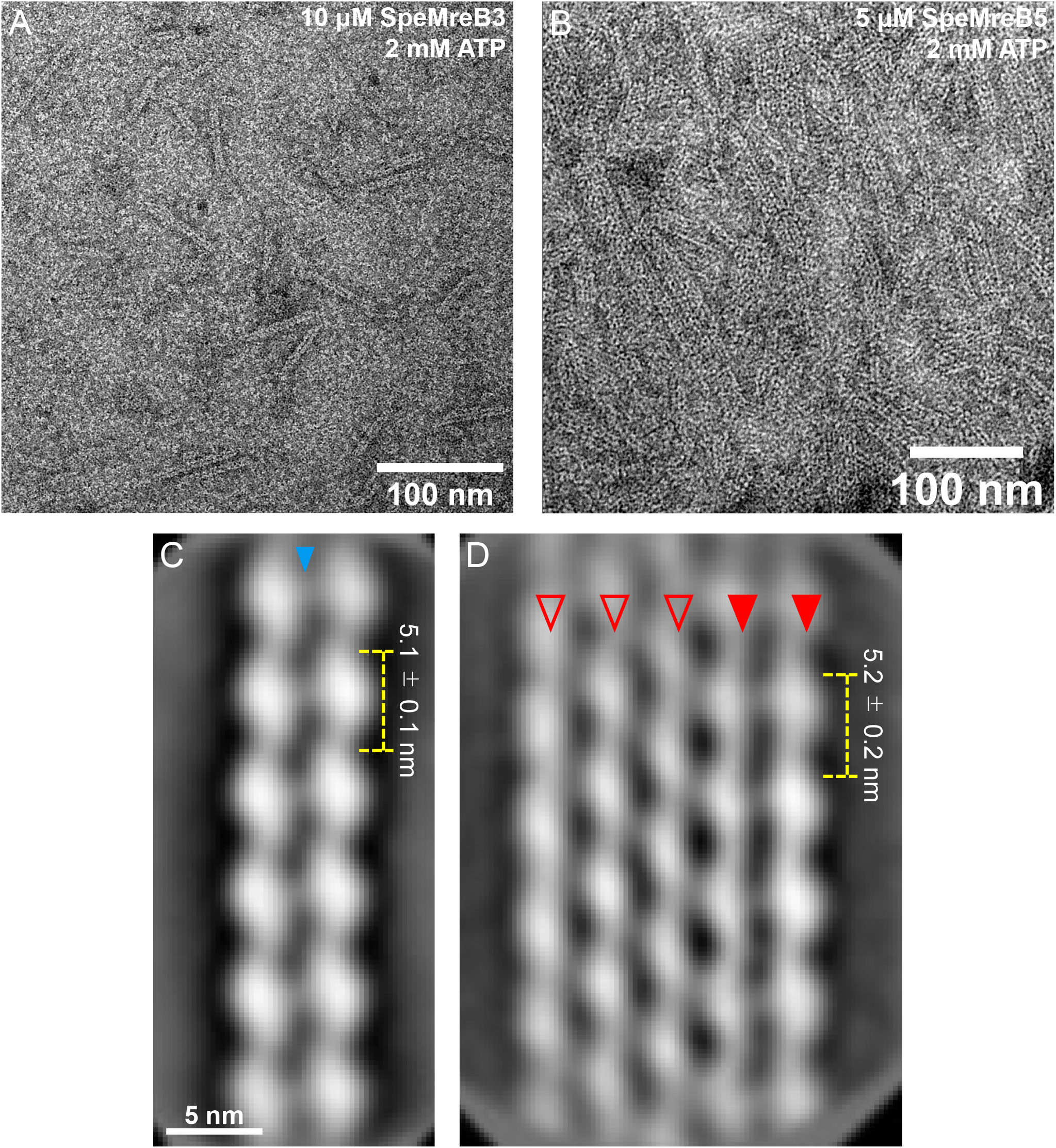
Structures of SpeMreB3 and SpeMreB5 filaments observed by EM. (**A-B**) Negative staining EM image of (**A**) 10 μM SpeMreB3 and (**B**) 5 μM SpeMreB5. (**C**) 2D averaged image of SpeMreB3 filaments averaged from 2874 particles. The estimated subunit repeats is 5.1 ± 0.1 nm. A weak electron density connecting the protofilaments is indicated by a triangle. (**D**) 2D averaged image of the five-stranded SpeMreB5 sheet structure averaged from 1,575 particles. The estimated subunit repeat is 5.2 ± 0.2 nm. The protofilaments in the antiparallel filaments and the other protofilaments are indicated by solid and open triangles, respectively.

To elucidate the subunit arrangement in these structures, we averaged the EM images using RELION ver. 3.1 or 4.0 (31). We picked 3,623 SpeMreB3 filaments and 117,740 SpeMreB5 sheet images. SpeMreB3 forms a double-stranded filament with a subunit repeat of 5.1 ± 0.1 nm (Fig. 1C). The two protofilaments were linked antiparallelly via weak density in a juxtaposed manner (Fig. 1C, Fig. S4A-C). These features resemble those of *Caulobacter crescentus* MreB (CcMreB) filaments (4). The SpeMreB5 sheet consists of variable numbers of parallelly aligning protofilaments with an antiparallel protofilament pair on one side of the sheet (Fig. 1D, Fig. S4D). Each parallel protofilament pair was organised in a staggered manner. As a result, the subunits were arranged diagonally over the parallel protofilaments in the sheet. The antiparallel protofilament was located at one edge of the sheet. The subunits in the antiparallel protofilament pair, as well as the SpeMreB3 filament, were arranged in a juxtaposed manner (Fig. 1C). Neither a sheet composed of parallel protofilament pairs nor one composed of only antiparallel protofilament pairs was observed. Moreover, none of the sheets had antiparallel double-stranded filaments on either side. These results indicate that the SpeMreB5 sheet is highly asymmetric and composed of two distinct inter-protofilament interactions.

A previous study reported that filament formation by *E. coli* MreB required nucleotide hydrolysis (5). To determine whether this is also the case for SpeMreB3 and SpeMreB5, we conducted negative staining EM observations of SpeMreB3 and SpeMreB5 incubated for 3 h in the presence of 2 mM Mg-AMPPNP or Mg-ADP. SpeMreB3 incubated with Mg-AMPPNP or Mg-ADP formed double-stranded filaments (Fig. S3C and D). SpeMreB5 with Mg-AMPPNP or Mg-ADP forms sheet structures (Fig. S3E and F). These results indicate that SpeMreB3 and SpeMreB5 polymerisations are driven by nucleotide binding rather than hydrolysis.

### Crystal structures of SpeMreB3

SpeMreB3 crystals suitable for X-ray experiments grew under several conditions but showed merohedral twinning. To overcome this problem, we methylated the lysine residues of SpeMreB3 and crystallised the methylated protein (Fig. S5A and B) (32, 33). The crystal structures of nucleotide-free (Nf) SpeMreB3 and its AMPPNP complex were determined at resolution of 1.90 and 1.75 Å, respectively (Fig. 2A, Fig. S6A, Table S1). SpeMreB3 adopts the canonical actin fold composed of four subdomains (IA, IB, IIA, and IIB) (34) and consists of the same secondary structure elements as in CcMreB, *Thremotoga maritima* (Tm) MreB, and SciMreB5, except for the very C-terminal region (3, 4, 35).

**Figure 2.**
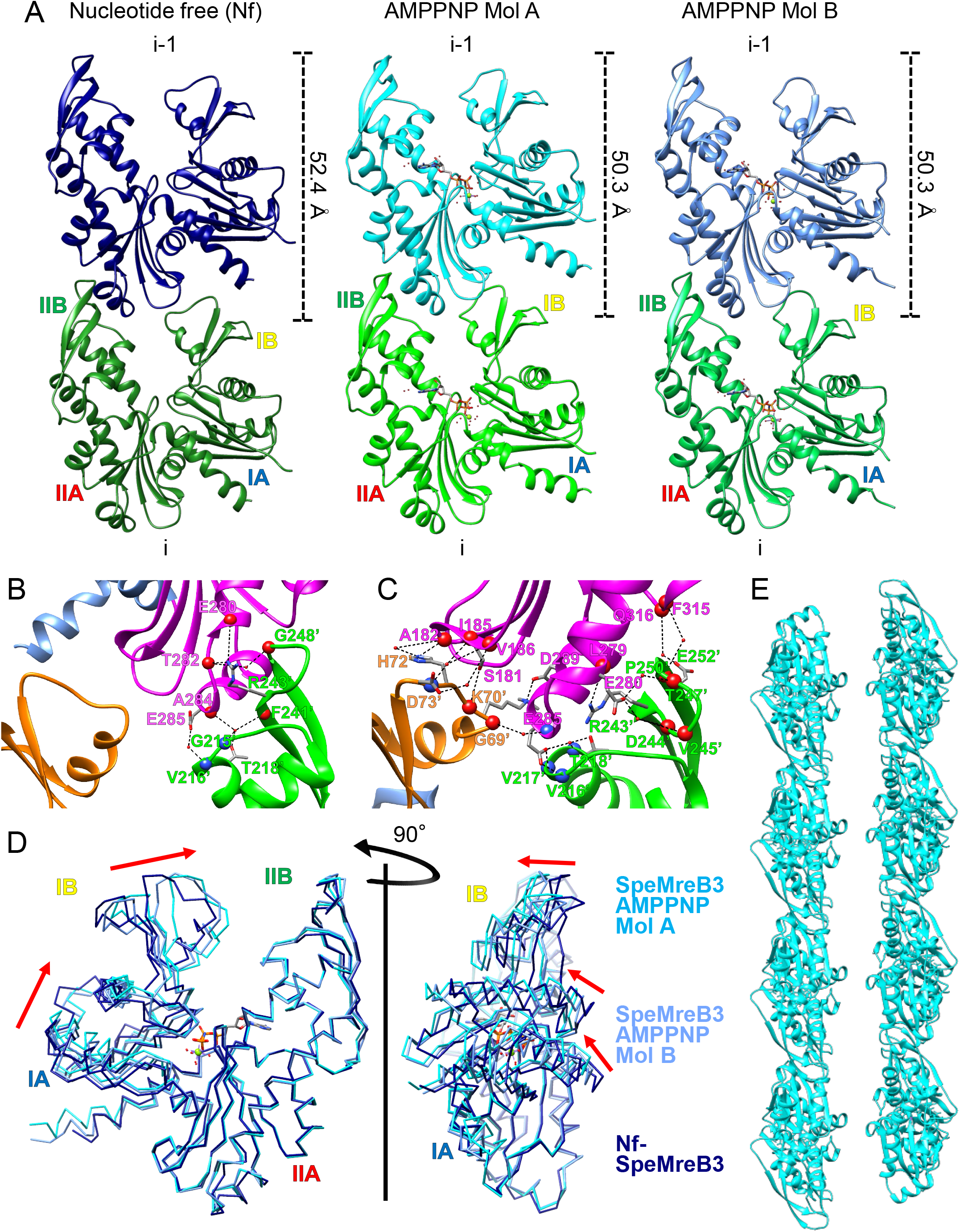
Crystal structures of SpeMreB3. (**A**) Protofilament structures in the crystals of the Nf-SpeMreB3 and SpeMreB3 AMPPNP complex. Two different conformations (Mol A and B) in the asymmetric unit of the SpeMreB3 AMPPNP complex crystal are shown in centre and right panels, respectively. Two subunits in the protofilaments are labelled as i and i-1. The subunit repeat is indicated at the right of each i-1 subunit. The four subdomains (IA, IB, IIA, and IIB) are labelled on i subunit. (**B-C**) Close up view of the subunit interface in the protofilament in the crystal of (**B**) Nf-SpeMreB3 and (**C**) the SpeMreB3 AMPPNP complex Mol A. Subdomains IA, IB, IIA, and IIB are indicated by ribbon representations coloured with pale blue, orange, magenta, and green, respectively. Hydrogen bonds and electrostatic interactions are indicated by broken lines. The residues involved in the hydrogen bonding or electrostatic interaction network are indicated by stick models or blue (backbone nitrogen atom) or red (backbone oxygen atom) spheres with labels. Water molecules involved in the interactions are shown with red small spheres. (**D**) Structural comparison of Mol A and B in the SpeMreB3 AMPPNP complex and Nf-SpeMreB3. The structures are superimposed to subdomains IIA and IIB of Mol A. Movement of the subdomain IA and IB is indicated by red arrows. (**E**) A ribbon representation of the filament structure of SpeMreB3 AMPPNP Mol A fit to the EM image (Fig. 1C, Fig. S4E).

The Nf-SpeMreB3 crystal belongs to the space group *P*2_1_ and contains a single molecule in an asymmetric unit. Nf-SpeMreB3 forms protofilaments along the crystal *a* axis in the *P*2_1_ crystal; thus, the protofilaments are arranged in an antiparallel manner. The subunit arrangement in the protofilament resembles that in the CcMreB, TmMreB, and SciMreB5 protofilaments in their respective crystals (Fig. 2A) (3, 4, 35). The interaction between IIA and IB ‘ (with and without a prime indicating i and i-1 subunits, respectively), which has been observed in the CcMreB, TmMreB, and SciMreB5 protofilaments, is not found in the Nf-SpeMreB3 protofilament, whereas the IIA-IIB ‘ interaction is conserved in Nf-SpeMreB3 (Fig. 2B). This IIA-IIB ‘ interaction is mediated by a hydrogen-bonding network and stabilised by an electrostatic interaction between E285 and the N-terminal end of the α-helix starting from V216 ‘.

The crystal of the SpeMreB3 AMPPNP complex includes two molecules (Mol-A and Mol-B) (Fig. 2A), which are related by a pseudo two-fold symmetry axis perpendicular to the crystal *a* axis, in an asymmetric unit. Each SpeMreB3 molecule in the asymmetric units forms a protofilament with those in the neighbouring unit cells along the crystal *a* axis. Thus, the SpeMreB3 AMPPNP complex crystal also contains antiparallel filaments, although their arrangement differs from that of the Nf-SpeMreB3 crystal. The two molecules show small differences in domain conformation, in which the nucleotide binding cleft in Mol-B is slightly wider than that in Mol-A (Fig. 2D, left). Compared with Nf-SpeMreB3, both clefts are narrower (Fig. 2D, left) and the overall conformation is more flattened (Fig. 2D, right). The IIA-IIB ‘ interaction area is wider than that of Nf-SpeMreB3 (Fig. 2B and C). As observed for Nf-SpeMreB3, E285 in the SpeMreB3 AMPPNP complex interacts electrostatically with the N-terminal end of the α-helix starting from V216 ‘. Similar to CcMreB, TmMreB, and SciMreB5 filaments (3, 4, 35), IIA in the SpeMreB3 AMPPNP complex also interacts with IB ‘, in which the interaction is mediated through a hydrogen-bonding network and stabilised by an electrostatic interaction between D289 and K70 ‘. The subunit interface area in the protofilament of SpeMreB3 AMPPNP is comparable to that of other MreB protofilaments, whereas that of Nf-SpeMreB3 is much smaller (Fig. S6B).

The subunit repeat along the protofilament in the crystal are in good agreement with that of the protofilament in the EM image (Fig. 1C, Fig. 2A). Therefore, we fitted the filament model in the crystal onto the 2D averaged EM image of the SpeMreB3 filament. A protofilament model of the SpeMreB3 AMPPNP complex fits well with the protofilament image (Fig. 2E, Fig. S4E), suggesting that the protofilament structure in the double-stranded filament (Fig. 1C) is the same as that in the crystal. However, none of the antiparallel filaments in the crystal fit the double-stranded filament in the EM image, indicating that the interaction stabilising the double-stranded filament in solution differs from that in the crystal.

### The “E ^…^ T - X - [D/E] “ motif is involved in the P_i_ release process of SpeMreB3 and SpeMreB5

Next, we measured the ATPase activity of SpeMreB3 and SpeMreB5 by P_i_ release assay. SpeMreB5 released P_i_ at a rate constant of 1.5 ± 0.2 nM (P_i_)/s/μM (protein) (Fig. 3A-C). However, the P_i_ release rate of SpeMreB3 was 33 times slower than that of SpeMreB5 (Fig. 3A and B).

**Figure 3.**
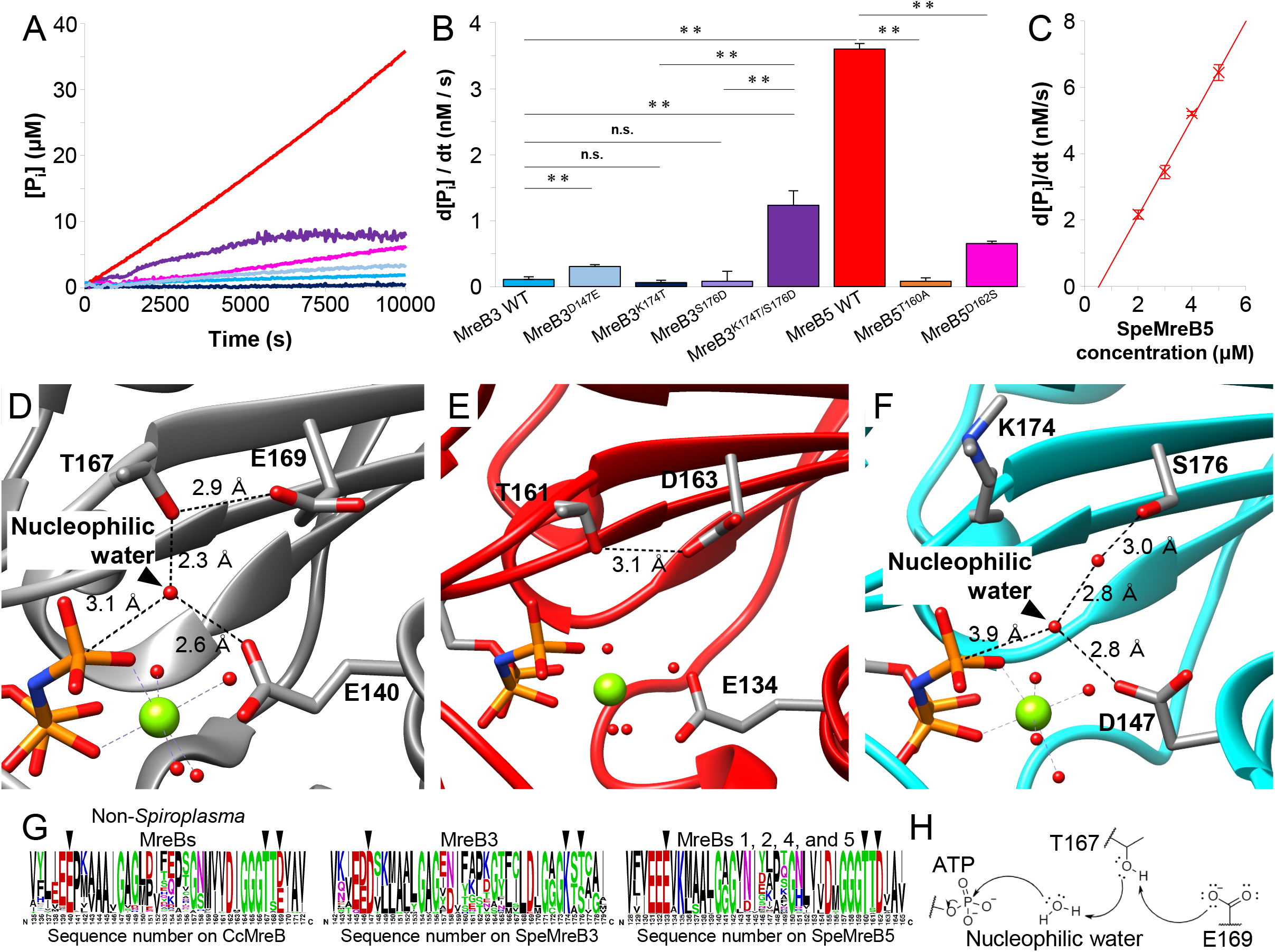
P_i_ release measurements and active site structures of SpeMreB3 and SpeMreB5. (**A**) Time course plots of P_i_ release of 3 μM SpeMreB3 wild type (cyan), SpeMreB3 D147E (pale blue), SpeMreB3 K174T (navy blue), SpeMreB3 K174T/S176D (purple), SpeMreB5 wild type (red), and SpeMreB5 D162S (pink) in the presence of 2 mM ATP. (**B**) P_i_ release rates of SpeMreBs estimated from panel A and Fig. S7G and H. Error bars indicate standard deviation (SD) from three repeated measurements. Symbols indicate *p*-value supported by Student ‘s *t*-test (**; *p* < 0.01, n.s.; *p* > 0.05). (**C**) Concentration dependence of the P_i_ release rates of SpeMreB5. Error bars indicate SD from three repeated measurements. (**D-F**) Close up view of the active sites of (**D**) the CcMreB AMPPNP complex (PDB: 4CZJ), (**E**) the SciMreB5 AMPPNP complex (PDB: 7BVY), and (**F**) the SpeMreB3 AMPPNP complex (Mol A). Mg^2+^ and water molecules are indicated as green and red spheres, respectively. (**G**) Weblogos of amino acid sequences around the ATP hydrolysis region on: (left) 4,832 MreB family proteins from non-*Spiroplasma* bacteria used in our previous study (9), (centre) 29 *Spiroplasma* MreB3, and (right) 171 *Spiroplasma* MreBs, except for MreB3. The corresponding amino acids for the core amino acids motif for ATP hydrolysis (E140, T167, and E169 in CcMreB) are indicated by triangles. (**H**) A working model for ATP hydrolysis in MreB family proteins. Residues corresponding to T167 and E169 in CcMreB, the nucleophilic water molecule, and γ-P_i_ of ATP are shown in this model. The unshared electron pairs of each atom on the residues and the water are indicated by two neighbouring dots. A putative electron transfer pathway is indicated by arrows.

To elucidate the structural basis of the slow P_i_ release rate, we compared the active site structures of CcMreB (PDB: 4CZJ), SciMreB5 (PDB: 7BVY), and SpeMreB3 in complex with AMPPNP (Fig. 3D-F). In the CcMreB structure, E140 and T167 coordinate with a putative nucleophilic water molecule that attacks the γ-P_i_ of ATP. T167 also forms a hydrogen bond with E169, which was thought to be unrelated to ATP hydrolysis (Fig. 3D) (4). These residues are structurally conserved in SciMreB5 (E134, T161, and D163 in Fig. 3E), although no nucleophilic water molecules were observed in the SciMreB5 structure. In the SpeMreB3 structure, D147 and K174 are located at positions corresponding to E140 and T167 in CcMreB, respectively. However, D147 is far from the putative nucleophilic water molecule and K174 does not interact with water molecules. The residue corresponding to E169 is replaced by serine (S176) in SpeMreB3, which does not interact with K174 (Fig. 3F). Therefore, the slow P_i_ release rate of SpeMreB3 could be attributed to these three residues.

To elucidate the role of these residues in the ATPase activity of SpeMreBs, we designed four mutant variants of SpeMreB3 (D147E, K174T, S176D, and K174T/S176D) and two variants of SpeMreB5 (T160A and D162S, which are equivalent to T161A and D163S mutations on SciMreB5, respectively (Fig. S1C)). All the mutant variants formed filamentous structures (Fig. S7A-F). The P_i_ release rate of SpeMreB3 D147E was three times higher than that of the wild type. SpeMreB3 K174T and SpeMreB3 S176D showed almost the same P_i_ release rate as the wild type, whereas SpeMreB3 K174T/S176D showed a 11-fold higher P_i_ release rate than the wild type (Fig. 3A and B, Fig. S7G). The P_i_ release rates of SpeMreB5 T160A and SpeMreB5 D162S were 50- and 5.6-fold slower than that of the wild type, respectively (Fig. 3A and B, Fig. S7H). These results indicate that the amino acid motif “E ^…^ T - X - [D/E] “ is important for the release of P_i_ from SpeMreBs, and that the Thr–Asp/Glu pair plays a distinct role from the first glutamate in the motif in ATPase activity.

To determine whether “E ^…^ T - X - [D/E] “ is conserved in the proteins of the MreB family, we analysed amino acid sequences of MreBs from all bacterial phyla (Fig. 3G). The motif is found to be conserved in 95.8% of MreB family proteins in non-*Spiroplasma* species and in 98.2% of *Spiroplasma* MreBs, except for MreB3 (Fig. 3G, left and right). The residues corresponding to E140 and T167 in CcMreB are replaced with aspartate and lysine, respectively, in all known *Spiroplasma* MreB3, as reported previously (9). Moreover, the residue corresponding to E169 in CcMreB is replaced with serine or threonine (Fig. 3G, centre). These findings suggest that the “E ^…^ T - X - [D/E] “ motif is important for the P_i_ release process of MreB family proteins, with the exception of *Spiroplasma* MreB3.

### Critical concentrations of SpeMreB3 and SpeMreB5 and their mutant variants

To evaluate the polymerisation activity of SpeMreB3 and SpeMreB5, we measured their critical concentrations, which reflect the minimum concentration required for polymerisation and the filament amounts at the steady state by sedimentation assay. However, significant amounts of proteins were precipitated even in the nucleotide-free buffer used for EM observation, while no filamentous structure was observed by EM (Fig. S3A and B, Fig. S8A), suggesting that amorphous aggregation affected the measurements. Therefore, we searched for a solution in which the proteins did not form aggregates without Mg-ATP, but polymerised in the presence of Mg-ATP. We found that a solution containing 20 mM Tris-HCl pH 8.0, 1 M NaCl, 200 mM L-arginine-HCl pH 8.0, and 5 mM DTT (buffer S) was suitable for the sedimentation assay (Fig. 4A and B, Fig. S3G and H).

**Figure 4.**
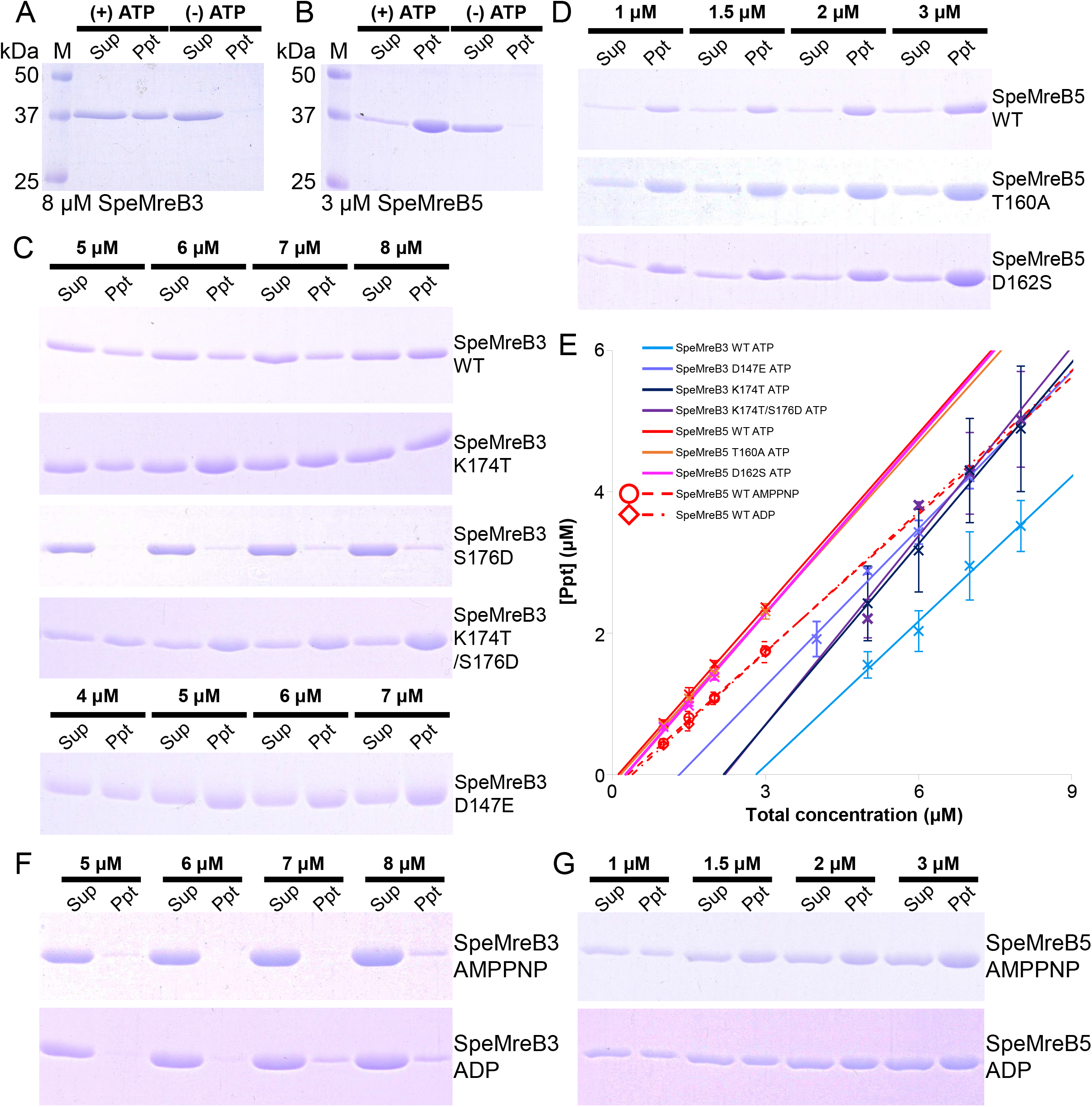
Sedimentation assay of the SpeMreBs. Each SpeMreB was incubated with the buffer S (20 mM Tris-HCl pH 8.0, 1 M NaCl, 200 mM Arginine-HCl pH 8.0, 5 mM DTT, 2 mM MgCl_2_, and 2 mM ATP) for 1 hour after initiating polymerization and subjected to the ultracentrifugation of 436,000 × *g*, 120 min at 23°C. Precipitates were resuspended with water equivalent amount to the samples. For SpeMreB3 and its mutant variants, each fraction was diluted three times before the preparation of the sample for SDS-PAGE. Each fraction was loaded onto 12.5% Laemmli gel and stained with Coomassie brilliant blue to quantify the concentrations. Fractions derived from a same sample were loaded onto adjacent lanes, and the total concentration of the sample is indicated on the lanes. (**A-B**) Sedimentation assay of (**A**) 8 μM SpeMreB3 and (**B**) 3 μM SpeMreB5 in the presence (left half lanes of each panel, (+) ATP) or absence (right half lanes of each panel, (-) ATP) of Mg-ATP. Protein size standards are visualized in Lane M with the molecular masses of each band on the left side. (**C-D**) Sedimentation assay over a range of concentrations of (**C**) SpeMreB3 and (**D**) SpeMreB5 wild type and mutant variants. (**E**) Quantified precipitation amounts of sedimented SpeMreBs. Precipitated SpeMreBs were resuspended in water equivalent amount to the sample, and the resulting concentrations of precipitated fractions were plotted over the total SpeMreB concentrations with linear fitting. Error bars indicate SD from five repeated measurements for SpeMreB5 wild type polymerized with ATP and ATP-analogs and three repeated measurements for the others. Critical concentrations were estimated as the x-intercept of each linear fit and summarized in Table 1. (**F-G**) Sedimentation assay over a range of concentrations of (**F**) SpeMreB3 and (**G**) SpeMreB5 polymerized with AMPPNP (top of each panel) or ADP (bottom of each panel) instead of ATP.

To estimate the time required to reach steady state, the protein samples in buffer S were incubated with 2 mM Mg-ATP for 1, 3, and 6 h and centrifuged. No significant difference was found in pellet amounts (Fig. S8B), indicating that polymerisation of SpeMreBs reached a steady state within 1 h. Therefore, the sedimentation assay was conducted after 1 h of incubation.

**Table 1.** Bulk critical concentration of SpeMreBs polymerized with the buffer S measured by the sedimentation assay. The values are indicated as mean ± SD from five repeating measurements for SpeMreB5 wild type polymerized with ATP and ATP-analogs and three repeated measurements for the others except for SpeMreB3 wild type polymerized with AMPPNP or ADP and SpeMreB3 S176D whose critical concentrations were estimated as the minimum concentrations where precipitates were detected. The row below the critical concentrations indicates *p*-value with the critical concentration of the represent WT polymerized with ATP supported by Student ‘s *t*-test. Symbols *, **, and n.s. indicate *p* < 0.05, *p* < 0.01, and *p* > 0.05, respectively.

|  | SpeMreB3<br>WT ATP | SpeMreB3<br>WT AMPPNP | SpeMreB3<br>WT ADP | SpeMreB3<br>D147E ATP | SpeMreB3<br>K174T ATP | SpeMreB3<br>S176D ATP | SpeMreB3<br>K174T/S176D<br>ATP | SpeMreB5<br>WT ATP | SpeMreB5<br>WT AMPPNP | SpeMreB5<br>WT ADP | SpeMreB5<br>T160A ATP | SpeMreB5<br>D162S ATP |
| --- | --- | --- | --- | --- | --- | --- | --- | --- | --- | --- | --- | --- |
| C<sub>C</sub> (μM) | 2.80 ± 0.34 | ≈ 8 | ≈ 7 | 1.26 ± 0.66 | 2.20 ± 0.50 | ≈ 6 | 2.15 ± 0.43 | 0.14 ± 0.07 | 0.29 ± 0.11 | 0.39 ± 0.09 | 0.15 ± 0.15 | 0.07 ± 0.03 |
| t - test<br>(v.s. WT ATP) | - | - | - | * (0.02) | n.s. (0.16) | - | n.s. (0.11) | - | * (0.03) | * * (0.001) | n.s. (0.87) | n.s. (0.20) |

The critical concentration of SpeMreB5 was estimated to be 0.14 ± 0.07 μM which is comparable to that of actin polymerized in a standard buffer for the actin polymerization, KMEI buffer (0.12-0.24 μM) and that of walled-bacterial MreBs under buffers akin to KMEI buffer (0.5 μM) (Fig. 4D and E, Table 1) (7, 8, 36, 37). In contrast, the critical concentration of SpeMreB3 was estimated to be 18 times higher than that of SpeMreB5 (Fig. 4C and E, Table 1).

Next, we determined the critical concentrations of the SpeMreB3 and SpeMreB5 mutant variants used in the P_i_ release assay. SpeMreB3 D147E showed a 2.2-fold lower critical concentration than SpeMreB3 wild type. SpeMreB3 S176D did not form sufficient precipitates to estimate the critical concentration using the linear fitting method. Therefore, we determined the minimum concentration at which precipitates were detected (6 μM) to be the approximate critical concentration of SpeMreB3 S176D (Table 1). These results suggest that these mutations affect the polymerisation activity of SpeMreB3. In contrast, the critical concentrations of the other mutant variants (SpeMreB3 K174T, SpeMreB3 K174T/S176D, SpeMreB5 T160A, and SpeMreB5 D162S) were not significantly different from those of their respective wild types (Fig. 4C-E, Table 1), suggesting that these point mutations do not affect the filament amounts of SpeMreB3 and SpeMreB5.

### ATP hydrolysis enhances the polymerization activity of SpeMreBs

To analyse the relationship between ATPase activity and polymerisation dynamics, we determined the critical concentrations of SpeMreB3 and SpeMreB5 in the presence of Mg-ADP or Mg-AMPPNP. The pellet amounts of SpeMreB3 polymerised with AMPPNP or ADP were less than those polymerised with ATP and too low to be applied to linear fitting (Fig. 4F). Therefore, the minimum concentrations forming precipitates were determined to be their approximate critical concentrations (Table 1). The critical concentration of SpeMreB5 was the lowest in the presence of ATP. It became 2- and 3-fold higher in the presence of AMP-PNP and ADP, respectively (Fig. 4E and G, Table 1). These results suggest that ATP hydrolysis enhances the polymerisation activities of SpeMreB3 and SpeMreB5.

## Discussion

### A model for polymerization dynamics of SpeMreBs

Electron microscopy and sedimentation assays of SpeMreB3 and SpeMreB5 showed that polymerisation was induced by nucleotide binding and enhanced by ATP hydrolysis (Fig. 1, Fig. 4, Fig. S3). Based on these results, we constructed a working model for the polymerisation dynamics of SpeMreB3 and SpeMreB5 (Fig. 5). First, SpeMreBs bind ATP and polymerise into filaments. Polymerised SpeMreBs hydrolyse ATP to stabilise the filaments. The SpeMreBs in the ADP-P_i_ state release phosphate after a certain duration to be converted to the ADP state, leading to depolymerisation. Finally, the depolymerised SpeMreBs in the ADP state replace ADP with ATP and return to the initial state in the polymerisation cycle.

**Figure 5.**
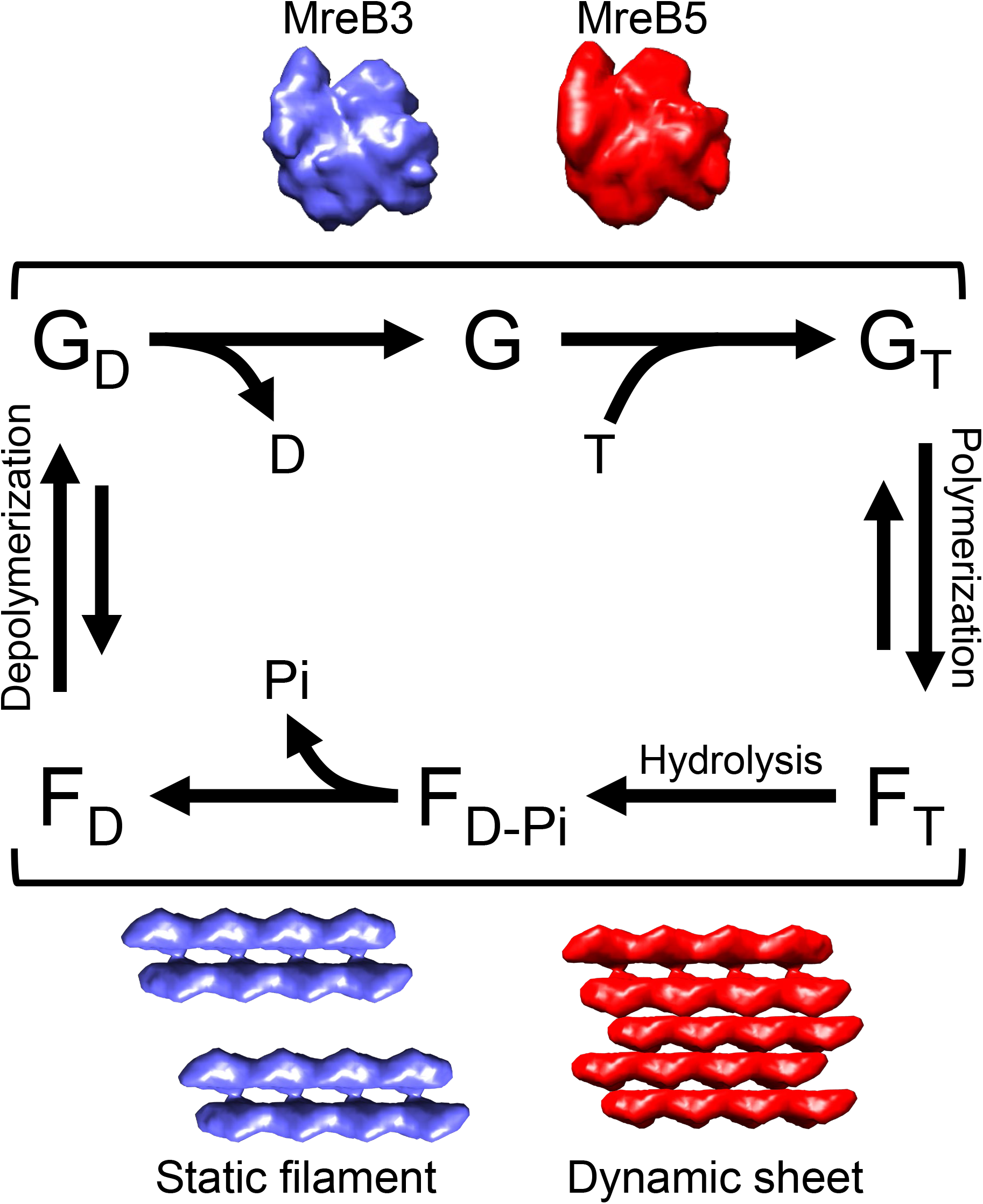
Working model of SpeMreB polymerization. ATP, ADP, and P_i_ are denoted as “T “, “D “, and “Pi “, respectively. The characters “G “ and “F “ indicate SpeMreBs in the monomeric and polymerized states, respectively, named as analogy based on actin. Bound nucleotides on SpeMreBs are indicated as a subscript. The schematic structures of the monomeric and polymerized SpeMreB3 and SpeMreB5 are indicated beside the corresponding positions on the polymerization cycles with the filament characters and the colours cyan and red, respectively.

This model is qualitatively applicable to both SpeMreB3 and SpeMreB5, although their rate constants differ. While our Nf-SpeMreB3 crystal structure forms a protofilament, we suppose that ATP binding is necessary to polymerise SpeMreB3 because filaments were not observed in the absence of nucleotides under solution conditions (Fig. 2A, Fig. 4A, Fig. S3A and B). Although some intra-protofilament interactions were observed in Nf-SpeMreB3 along the filament axis, the interaction area is smaller than that in SpeMreB3-AMPPNP (Fig. 2B and C, Fig. S6B). This difference is derived from domain closure upon AMPPNP binding (Fig. 2D), suggesting that this conformational change is necessary to form filaments in solution. Previous studies using CcMreB and TmMreB showed that domain closure is induced upon polymerisation (4, 38), whereas our structural analyses showed that nucleotide binding is also essential for domain closure (Fig. 2D).

The working model was qualitatively consistent with that of other actin superfamily proteins (30). However, the equilibrium balance of polymerisation dynamics is likely to be uncommon. The critical concentration of actin polymerised with ADP was 18-fold higher than that polymerised with ATP (37, 39). On the other hand, the critical concentrations of SpeMreBs polymerised with ADP were only approximately two times higher than those with ATP (Fig. 4E-G, Table 1), suggesting that SpeMreB3 and SpeMreB5 filaments remain stable after P_i_ release, unlike actin filaments. Structural differences between actin and SpeMreBs may cause differences in their critical concentrations. In the actin filament, the D-loop in subdomain IB (subdomain 2 in the nomenclature of actin) forms a major intra-protofilament interaction that is attenuated upon P_i_ release (40). SpeMreB3 and SpeMreB5 lack a loop corresponding to the D-loop of actin, as well as MreB of walled bacteria, and their intra-protofilament interaction via subdomain IB is less than that of actin (Fig. 2C) (3, 4, 35). These facts suggest that subunit turnover rates of SpeMreBs are slower than those of actin. This may be a fundamental feature of MreB family proteins, as the critical concentration of TmMreB polymerised with ADP is approximately 2-fold higher than that polymerised with ATP, as well as SpeMreB3 and SpeMreB5 (8).

### ATP hydrolysis mechanism of MreB family proteins

Through our P_i_ release assays of SpeMreB3 and SpeMreB5 (Fig. 3, Fig. S7G and H), we determined two players of ATPase activity, the threonine–acidic residue pair and the conserved glutamate corresponding to E140 in CcMreB. These residues form a hydrogen-bonding network with a putative nucleophilic water molecule in the CcMreB structure (Fig. 3D), suggesting that these residues play a role in ATP hydrolysis. A previous study showed that E140 and T167 in CcMreB align nucleophilic water to an appropriate position for ATP hydrolysis (4). However, it was not clear which residue plays a role in proton elimination from the water molecule, which is a necessary step for ATP hydrolysis. Based on our findings, we updated the role of active-site residues in ATP hydrolysis using CcMreB as a model (Fig. 3H). E169 eliminates the proton of the side chain hydroxy group of T167 to activate it as an acidic catalyst for proton elimination of nucleophilic water. E140 adjusts the position of the nucleophilic water to be suitable for attacking the γ-P_i_ of ATP (Fig. 3D). This reaction mechanism is similar to that proposed for skeletal actin, where Q137, corresponding to E140 in CcMreB, is responsible for the positioning of nucleophilic water, and H161, corresponding to E169 in CcMreB, is responsible for the proton elimination of nucleophilic water (40). The amino acid motif “E ^…^ T - X - [DE] “ is mostly conserved in the MreB family proteins, except for *Spiroplasma* MreB3 (Fig. 3G), suggesting that the ATP hydrolysis mechanism proposed here is conserved in most MreB family proteins.

### Role of the polymerization characteristics of SpeMreB3 and SpeMreB5 for *Spiroplasma* swimming

A previous study on *S. poulsonii* MreBs using a heterologous expression system suggested that MreB3 and MreB5 play distinct roles in the cell (26). Our results showed that SpeMreB3 and SpeMreB5 polymerise into different filamentous structures with distinct ATPase activities and critical concentrations (Fig. 1, Fig. 3, Fig. 4). These differences in polymerisation characteristics may be related to the differences in the roles of SpeMreB3 and SpeMreB5. MreB5 has been suggested as an actuator that changes the conformation of the internal ribbon structure to drive the cell (26). Our EM observations showed that SpeMreB5 forms sheet structures with two patterns of inter-protofilament interactions (Fig. 1D), which have not been observed in walled-bacterial MreBs. Because of this distinct interaction patterns, the sheet contained both parallel and antiparallel alignments of the two protofilaments. The parallel alignment can lead to anisotropy in the overall sheet structure which meets the requirement to cause directional movement of *Spiroplasma* swimming. MreB3 is suggested to anchor MreB1 and/or MreB4 filaments to the membrane to form a fixed structure (26). Therefore, this role does not require filament polarity. SpeMreB3 forms a double-stranded filament with antiparallel polarity (Fig. 1C), which is consistent with this requirement.

SpeMreB3 showed low ATPase activity (Fig. 3A and B, Fig. S7G). To the best of our knowledge, this is one of the lowest activities of the actin superfamily proteins. The amino acid residues responsible for the low ATPase activity are conserved in MreB3 (Fig. 3G, centre). These findings suggest that low ATPase activity is also involved in SpeMreB3 function. As subunit turnover requires the completion of the ATPase cycle (Fig. 5), a low ATPase activity leads to a slow turnover of subunits, suggesting that the SpeMreB3 filament is stable and may stably anchor the SpeMreB1 and/or SpeMreB4 filaments onto the membrane. This is consistent with a previous study showing that *S. poulsonii* MreB3 forms static filaments in a heterologous expression system (26).

### Effect of SpeMreB3 methylation on polymerization

Lysine residues were methylated to obtain SpeMreB3 crystals suitable for structural determination (Fig. S5A and B). Methylated SpeMreB3 was polymerised, as observed in unmethylated filaments, when the concentration was approximately 10-fold higher (Fig. S5C). Nevertheless, in the sedimentation assay, pellets containing 8 μM methylated SpeMreB3 were undetectable (Fig. S5D), indicating that the polymerisation activity was attenuated by methylation. One cause may be the inhibition of lysine residue-mediated interactions involved in filament formation. Our crystal structure of the SpeMreB3 AMPPNP complex showed that K70, which is involved in the intra-protofilament interaction, was di-methylated (Fig. 2C). In methylated SpeMreB3, this interaction may be disturbed in solution conditions, most likely due to a decrease in the degree of freedom of the interaction. In the structure of SpeMreB3 AMPPNP complex, K174 is di-methylated. We cannot rule out the possibility that this methylation changes the position of the residue, preventing interaction with the nucleophilic water (Fig. 3F) and slowing the ATP hydrolysis rate, as suggested by the decrease in the pellet amount in the sedimentation assay (Fig. 3F, Fig. S5D). However, this methylation is unlikely to affect the above discussion on ATP hydrolysis because the P_i_ release rate was not changed by introducing the single K174T mutation (Fig. 3A, B, and F, Fig. S7B).

## Materials and Methods

### SpeMreBs cloning and expression

The DNA sequences of SpeMreB3 and SpeMreB5 (23) were codon optimized for *E. coli* expression and synthesized as the fusion with pUC57 (GenScript, Piscataway, NJ). The DNA fragments encoding SpeMreB3 and SpeMreB5 were excised using restriction enzymes NdeI and BamHI and inserted into pET-15b (Novagen, Madison, WI) or pCold-15b, which was constructed from pCold I (Takara Bio Inc., Kusatsu, Japan) by replacing the sequences of the histidine tag and proteinase digestion site with that of pET-15b. Each construct was transformed into *E. coli* BL21 (DE3) and C43 (DE3) cells. *E. coli* carrying the constructed plasmid were grown overnight in LB medium in the presence of 50 μg/mL ampicillin at 37°C. The culture was diluted with fresh medium and cultivated at 37°C. When the OD_600_ value reached 0.4-0.6, IPTG was added to a final concentration of 1 mM and the cultures were incubated for 24 h at 15°C. The cells were harvested, washed twice with PBS (10 mM Na_2_HPO_4_, 2 mM NaH_2_PO_4_, 3 mM KCl, and 137 mM NaCl), and stored at L80°C until further use.

### SpeMreB purification

The wild type and mutant variants of SpeMreB3 were purified as the fusion with a 6×histidine-tag at the N-terminus as follows. The cell pellets harvested from 1-L cultures were resuspended in 20-40 mL buffer A (50 mM Tris-HCl pH 8.0 at 25°C, 300 mM NaCl, 50 mM imidazole-HCl pH 8.0 at 25°C) and sonicated with a probe sonicator (Nissei, Ultrasonic Homogeniser). The cell lysate was then centrifuged (100,000 × *g*, at 4°C, 30 min). The supernatant was subjected to HisTrap HP column (Cytiva, Wauwatosa, WI, USA), washed with 10 column volumes of buffer A, and eluted with 13 mL of buffer A containing 230 mM imidazole-HCl pH 8.0 at 25°C. The eluted SpeMreBs were further subjected to a HiLoad 26/600 Superdex 200 pg column (Cytiva) at 4°C equilibrated with buffer B (20 mM Tris-HCl pH 8.0 at 25°C, 300 mM NaCl). For SpeMreB5 and its mutant variants, the magnitude of centrifugation was decreased to 12,000 × *g*. For the purification of SpeMreB3 for the crystallisation experiments, buffer C (10 mM Tris-HCl pH 8.0 at 25°C and 150 mM NaCl) was used for gel filtration with a HiLoad 26/600 Superdex 200 pg column instead of buffer B. The protein concentrations were determined from the absorbance at 280 nm measured using NanoDrop One (Thermo Fisher Scientific) with the following absorption coefficients: 0.474 (mg/mL)^-1^cm^-1^ for SpeMreB3 and its mutant variants and 0.578 (mg/mL) ^-1^cm^-1^ for SpeMreB5 and its mutant variants.

### Methylation of SpeMreB3

SpeMreB3 eluted from Ni^2+^-NTA chromatography was subjected to HiPrep 26/10 Desalting (Cytiva) equilibrated with buffer D (50 mM HEPES-NaOH pH 7.5 at 25°C and 250 mM NaCl) or dialysed with buffer D to replace the buffer. To methylate the lysine residues of SpeMreB3, dimethylamine-borane complex (DMAB) (Merck) and formaldehyde (Merck) were added to final concentrations of 20 and 40 mM, respectively (32, 33), and the sample was incubated for 2 h at 4°C. The methylated sample was subjected to HiPrep 26/10 Desalting equilibrated with buffer D to remove excess DMAB and formaldehyde, concentrated to less than 13 mL using Amicon Ultra 10 K (Merck), and subjected to gel filtration chromatography on a HiLoad 26/600 Superdex 200 pg. The column was equilibrated with buffer B at 4°C. The methylation ratio of methylated SpeMreB3 reached 96.5 ± 2.2%, as confirmed by MALDI-TOF mass spectrometry (Fig. S5B). To prepare methylated SpeMreB3 for crystallisation, the following three steps were modified from the method described above: (1) After Ni^2+^-NTA affinity chromatography, the mixture of the sample and 100 units of thrombin (Cytiva) was dialysed overnight at 4°C in buffer D to cleave the histidine tag from SpeMreB3; (2) After 2 h of incubation with 20 mM DMAB and 40 mM formaldehyde, two incubation steps were added before the removal of excess DMAB and formaldehyde. DMAB and formaldehyde were added at concentrations of 40 and 80 mM, respectively, and the sample was additionally incubated for 2 h at 4°C. DMAB was further added to 50 mM, and the sample was incubated overnight at 4°C. In this procedure, although the terminal 7-13 amino acids of methylated SpeMreB3 were cleaved, and small amounts of degraded products appeared even after the final purification step, methylated SpeMreB3 was successfully crystallised; (3) Buffer C was used for gel filtration on a HiLoad 26/600 Superdex 200 pg column, instead of buffer B.

### SpeMreB polymerization

For the preparation of samples for EM observations and the measurement of P_i_ release rates, SpeMreBs were polymerised in a standard buffer (20 mM Tris-HCl pH 7.5 at 25°C, 100 mM KCl, 5 mM DTT, 2 mM MgCl_2_, and 2 mM ATP). For the sedimentation assay, SpeMreBs were polymerised in buffer S (20 mM Tris-HCl pH 8.0 at 25°C, 1 M NaCl, 200 mM L-arginine-HCl pH 8.0, 5 mM DTT, 2 mM MgCl_2_, and 2 mM ATP). Prior to polymerisation, the SpeMreBs buffer was changed from buffer B to the desired buffer in the absence of DTT, MgCl_2_, and ATP. Monomeric SpeMreBs with a concentration lower than the desired concentration were concentrated using an Amicon Ultra 10 K to desired. Samples were centrifuged to remove aggregates and DTT, MgCl_2_, and ATP were added to initiate polymerisation. All the polymerisation reactions were performed at room temperature (∼25°C).

### Electron microscopy (EM)

A sample (4 μL) was placed onto a 400-mesh copper grid coated with carbon for 1 min at room temperature, washed with 10 μL of water, stained for 45 s with 2% (w/v) uranyl acetate, air-dried, and observed under a JEOL JEM-1010 transmission electron microscope (Tokyo, Japan) at 80 kV, equipped with a FastScan-F214T charge-coupled device camera (TVIPS, Gauting, Germany). To obtain SpeMreB3 and SpeMreB5 images for 2D averaging, 10 μM SpeMreB3 and 5 μM SpeMreB5 polymerised with standard buffer were diluted to 3 μM immediately before sample placement onto a grid. For image averaging, SpeMreB images were automatically picked using RELION ver. 3.1 (31) as helical objects, and they were segmented in a box of 128 × 128 pixels with 90% overlap. These images were subjected to estimation of the contrast transfer function and reference-free 2D class averaging using RELION ver. 3.1 or 4.0 (31). The subunit repeat in the 2D averaged images were estimated as the distances between the minimal values of grayscale profiles quantified using ImageJ (National Institutes of Health; http://rsb.info.nih.gov/ij/).

### Crystallization and structural determination

Crystallisation screening was performed using the sitting-drop vapor-diffusion technique with the following screening kits: Wizard Classic I-II (Rigaku Reagents, Inc., Bainbidge Island, USA), Wizard Cryo I-II (Rigaku Reagents, Inc.), PEG/Ion Screen I-II (Hampton Research, Alison Viejo, USA), Crystal Screen I-II (Hampton Research), SaltRx I-II (Hampton Research), and PEG/Ion 400 (Hampton Research), at 4°C and 20°C. Crystals of Nf-SpeMreB3 used for X-ray data collection were grown at 4°C from drops prepared by mixing 0.5 μL of protein solution (5 mg/mL) in buffer C with an equivalent volume of reservoir solution containing 100 mM MES-NaOH pH 6.0, 20% (w/v) PEG-8000, and 200 mM calcium acetate. The crystals belong to the space group *P*2_1_ with unit cell dimensions of *a* = 52.4, *b* = 68.1, *c* = 54.6 Å, and β = 91.7°. The SpeMreB3 AMPPNP complex crystals used for X-ray data collection were obtained at 20°C from drops prepared by mixing 0.5 μL of protein solution (5 mg/mL) in buffer C containing 5 mM MgCl_2_ and 5 mM Li_4_AMPPNP with the equivalent volume of reservoir solution containing 100 mM Acetate-NaOH pH 4.6, 30% (w/v) PEG-4000, and 200 mM ammonium acetate. The crystals belong to the space group *P*2_1_ with unit cell dimensions of *a* = 50.3, *b* = 56.3, *c* = 120.5 Å, and β = 90.6°.

X-ray diffraction data were measured at synchrotron beamlines BL41XU and BL45XU at SPring-8 (Harima, Japan) with the approval of the Japan Synchrotron Radiation Research Institute (JASRI) (proposal no. 2018A2567, 2018B2567, 2019A2550, and 2019B2550). The crystals were soaked in a 1:9 mixture of glycerol and reservoir solution for several seconds before being transferred to liquid nitrogen for freezing. The X-ray diffraction data were measured at 100 K under nitrogen gas flow. The diffraction data were processed using MOSFLM (41) and scaled using Aimless (42). The initial phase was determined by molecular replacement (MR) with Phaser (43) using the CcMreB structure (PDB ID: 4CZL) as a search model. An atomic model of Nf-SpeMreB3 was constructed using Coot (44) and refined using Phenix (43). The final model includes P17–L344. The refined structure of Nf-SpeMreB3 was used as a search model for molecular replacement of the SpeMreB3 AMPPNP complex. An atomic model of the SpeMreB3 AMPPNP complex was built using Coot (44) and refined using Phenix (43). The final model includes P17–N348 for Mol-A and P18–E347 for Mol-B. The data collection and refinement statistics are summarised in Table S1.

### P_i_ release assay

P_i_ release was measured using the EnzChek kit (Thermo Fisher Scientific), in which P_i_ concentration is traced by the absorbance at 360 nm (A_360_) of the reaction product between P_i_ and 2-amino-6-mercapto-7-methylpurine riboside (MESG), a molecular probe for P_i_ (8, 45-47). Reactions were initiated by adding a mixture of MgCl_2_, ATP, and MESG to 3 μM SpeMreBs in standard buffer without Mg-ATP. The A_360_ values of SpeMreBs without MESG and with the buffer containing MESG were subtracted from those measured for SpeMreBs with MESG. Data were collected using a Varioskan Flash (Thermo Scientific), which has an initial measurement delay of approximately 1 min.

### Sedimentation assay

SpeMreBs with a volume of 200 μL were polymerised for 1 to 6 h and centrifuged (100,000 rpm, 23°C, 120 min) in a TLA-100 rotor (Beckman Coulter). The pellet was resuspended in 200 μL of water. The supernatant and pellet fractions were subjected to electrophoresis on a 12.5% Laemmli gel and stained with Coomassie Brilliant Blue R-250 to analyse the concentration of each fraction. The band intensities of SpeMreBs were quantified using the ImageJ software. The critical concentration was determined as the x-intercept of a linear fit with the precipitate amounts at the steady state of polymerisation over the total SpeMreB concentrations.

## Supporting information

MreBpolymerizationPaper_20220324_bioRxiv

## Acknowledgments

We thank Ms. Junko Shiomi and Ms. Tomomi Shimonaka for technical assistance with SpeMreB expression and MALDI-TOF MASS spectrometry, respectively. We also thank Dr. Yoshihiro Yamaguchi for providing *E. coli* C43 (DE3), Mr. Yuhei O Tahara, and Dr. Takuma Toyonaga for technical assistance with electron microscopy and its image analyses (all of which are affiliated with the Graduate School of Science, Osaka City University), Dr. Norihiro Takekawa, and Mr. Motoshi Sakai (Graduate School of Science, Osaka University) for technical assistance with the data collection of X-ray diffraction experiments, and Dr. Miki Kinoshita (Graduate School of Frontier Bioscience, Osaka University) for providing reagents for SpeMreB3 methylation. The English language of the manuscript was confirmed by Dr. Timothy Day (Graduate School of Frontier Bioscience, Osaka University). This study was supported by Grants-in-Aid for Scientific Research (A and C) (MEXT KAKENHI) (grant number JP17H01544 to MM and JP20K06591 to IF), JST CREST (grant number JPMJCR19S5 to MM), the Research Foundation of Opto-Science and Technology to IF, and the Osaka City University (OCU) Strategic Research Grant 2019 to IF. DT is a recipient of a Japanese scholarship from JEES Kureha (Toyobo) and the Ono Scholarship Foundation.

## Author Contributions

The experimental design was prepared by DT, IF, and MM. Data correction of the X-ray diffraction experiments and structural determination were performed by KI. EM image analyses were performed by DT, AN, and YS. All other experiments and data analyses were performed by DT. The manuscript was written by DT, IF, and MM. All authors approved the final version of the manuscript.

