## Supplementary material for "ATP dependent polymerization dynamics of bacterial actin proteins involved in *Spiroplasma* swimming": MreBpolymerizationPaper_20220324_bioRxiv

**Table S1. X-ray data collection and refinement statistics.** Values in parentheses are for the highest resolution shell. $R_w = \sum || F_o | - | F_c || / \sum | F_o |$, $R_{free} = \sum || F_o | - | F_c || / \sum | F_o |$.

**Figure S1. Sequence and structural characters of SpeMreBs.** (**A**) Summary of sequence identities between two SpeMreBs for the all combinations. Each combination is connected with a black line in which the darkness varies corresponding to the sequence identity shown beside the line. (**B-C**) Sequence comparisons (**B**) among SpeMreBs and walled-bacterial MreBs of *Thermotoga maritima* (Tm), *Caulobacter crescentus* (Cc), *Escherichia coli* (Ec), and *Bacillus subtilis* (Bs) MreBs, and (**C**) between SciMreB5 and SpeMreB5. Subdomains IA, IB, IIA, and IIB are indicated by half parentheses colored with blue, orange, red, and green, respectively. Nucleotide binding cleft, nucleotide hydrolysis, and N-terminal amphipathic helices are surrounded by orange-, cyan-, and purple-colored boxes, respectively. Asterisks (*), colons (:), and periods (.) below each amino acid indicate that the amino acids are identical, strongly similar (scoring greater than 0.5 in the Gonnet PAM 250 matrix), and weakly similar (scoring greater than 0.0 but no more than 0.5 in the Gonnet PAM 250 matrix), respectively.

**Figure S2. Purification and gel filtration profile of SpeMreB3 and SpeMreB5. (A-B)** Purification procedure of (**A**) SpeMreB3 and (**B**) SpeMreB5 from *E. coli* BL21 (DE3) carrying a codon optimized *spemreB* gene fused with pET-15b. The following fractions during the procedure are visualized by Coomassie stained 12.5% Laemmli gel: (1) Fraction 1: Lysate of *E. coli* carrying no plasmids; (2) Fraction 2: Lysate of IPTG uninduced *E. coli* carrying a plasmid for SpeMreB expression; (3) Fraction 3: Lysate of *E. coli* expressing an SpeMreB by IPTG induction; (4) Fraction 4 and 5: Soluble (4) and insoluble (5) fractions of an SpeMreB expressing *E. coli* lysate; (5) Fraction 6 and 7: Flow through (6) and elution (7) fractions of $Ni^{2+}$ affinity chromatography; (6) Fraction 8: A peak fraction of gel filtration. Protein size standards are visualized in Lane M with the molecular masses of each band on the right side. Triangles indicate the regions where histidine-tag fused SpeMreB bands appear. (**C-D**) Gel filtration profile of (**C**) SpeMreB3 and (**D**) SpeMreB5 with HiLoad 26/600 Superdex 200 pg. (Cytiva). Triangles indicate the peaks of the SpeMreB elution fractions. The elution volume of bovine γ-globulin (158 kDa), chicken ovalbumin (44 kDa), and horse myoglobin (17 kDa) are plotted as diamonds over the log of their molecular masses. From these elution spectra, the molecular masses of histidine-tag fused SpeMreB3 and SpeMreB5 were estimated to be 44 and 46 kDa, respectively, indicating that both SpeMreBs are purified as monomers.

**Figure S3. Structures of SpeMreB3 and SpeMreB5 filaments under various conditions.** (**A-B**) Negative staining EM images of (**A**) 10 μM SpeMreB3 and (**B**) 5 μM SpeMreB5 in the absence of Mg-ATP. (**C-D**) Negative staining EM images of 10 μM SpeMreB3 polymerized with (**C**) 2 mM Mg-AMPPNP and (**D**) 2 mM Mg-ADP instead of Mg-ATP. (**E-F**) Negative staining EM image of 5 μM SpeMreB5 polymerized with (**E**) 2 mM Mg-AMPPNP and (**F**) 2 mM Mg-ADP instead of Mg-ATP. (**G-H**) Negative staining EM image of (**G**) 10 μM SpeMreB3 and (**H**) 5 μM SpeMreB5 polymerized in buffer S.

**Figure S4. Structural characteristics of SpeMreB3 and SpeMreB5 filaments.** (**A**) Processed 2D averaged images of the SpeMreB3 filament. The original image is shown in the centre. The horizontally flipped, and 180° rotated images are shown on the left and right sides, respectively. Blue arrows in each SpeMreB3 subunit point from the swollen part at the outside to the end of a subunit at the opposite side. The topology of the original

SpeMreB3 filament is the same as that of the 180° rotated filament rather than the horizontally flipped filament, indicating the antiparallel polarity. (**B**-**C**) Schematic illustrations of non-helical filaments with (**B**) antiparallel and (**C**) parallel polarity. Both filaments show $C_2$-symmetry at an axis indicated as orange bars. (**D**) SpeMreB5 sheet images composed of three (left) and four (right) protofilaments. The subunit repeat of the protofilaments and the particle number of triple- and quadruple-stranded sheets are 5.2 ± 0.2 and 5.2 ± 0.3 nm, respectively, and 1,864 and 643, respectively. The protofilaments in the antiparallel filaments and the other protofilaments are indicated by solid and open triangles, respectively. (**E**) Fitted ribbon represented SpeMreB3 protofilaments composed of Mol A of AMPPNP complex into the 2D averaged EM image.

**Figure S5. Purification and polymerization assay of methylated SpeMreB3.** (**A**) Purification procedure of methylated SpeMreB3 from *E. coli* C43 (DE3) carrying pCold-15b fused with a gene to express SpeMreB3. The following fractions during the procedure are visualized by Coomassie stained 12.5% Laemmli gel: (1) Fraction 1: Lysate of *E. coli* expressing SpeMreB3 by IPTG induction; (2) Fraction 2 and 3: Soluble (2) and insoluble (3) fractions of SpeMreB3 expressing *E. coli* lysate; (3) Fraction 4 and 5: Flow through (4) and elution (5) fractions of $Ni^{2+}$ affinity chromatography; (4) Fraction 6: SpeMreB3 after the methylation treatment; (5) Fraction 7: Load fraction for gel filtration; (6) Others: Elution fractions of gel filtration. Protein size standards are visualized in Lane M with the molecular masses of each band on the left side. Cyan and black triangles indicate the regions for unmethylated SpeMreB3 and methylated SpeMreB3, respectively. (**B**) Mass spectra of (cyan) unmethylated SpeMreB3 and (black) methylated SpeMreB3. From these spectra, the increase of SpeMreB3 molecular mass by methylation is calculated as 891 ± 20 Da; namely, the estimated methylating ratio is 96.5 ± 2.2% because SpeMreB3 has 33 lysine residues and the molecular mass is increased 28 Da per lysine by the reaction. (**C**)

Negative staining EM image of 75 μM methylated SpeMreB3 polymerized in the standard buffer for 3 hours. (**D**) Sedimentation assay of 8 μM methylated SpeMreB3. It was polymerized for 3 hours with the buffer S and subjected to the centrifugation of 436,000 × *g* for 120 min at 23°C. Each fraction was diluted three times before the preparation of the sample for SDS-PAGE. Protein size standards are visualized in the lane M with the molecular masses of each band on the right side.

**Figure S6. Electron density map of the active site of SpeMreB3 and the subunit interface in protofilaments of MreBs.** (**A**) Nucleotide binding site. The 2Fo-Fc density map contoured at 2.0 sigma (blue) and the Fo-Fc density map contoured at 2.0 sigma (green) are superimposed onto the refined model. The AMPPNP-Mg moiety was omitted from the model for phase calculation. (**B**) Surface representation of two subunits in the protofilaments of SpeMreB3 AMPPNP Mol A, Nf-SpeMreB3, CcMreB (PDB: 4CZJ), TmMreB (PDB: 1JCG), and SciMreB5 (PDB: 7BVY). The four subdomains (IA, IB, IIA, and IIB) of i subunit are labeled. The subunit interface is colored with red. The subunit interface area with 2.5 Å cut off (calculated using Chimera 1.13.1) is shown below each model.

**Figure S7. Filament structures and $P_i$ release measurements of mutant variants of SpeMreB3 and SpeMreB5.** (**A-F**) Negative staining EM images of (**A**) 10 μM SpeMreB3 D147E, (**B**) 10 μM SpeMreB3 K174T, (**C**) 10 μM SpeMreB3 S176D, (**D**) 10 μM SpeMreB3 K174T/S176D, (**E**) 5 μM SpeMreB5 T160A, and (**F**) 5 μM SpeMreB5 D162S polymerized in the standard buffer for 3 hours. (**G**) Time course plots of $P_i$ release of 3 μM SpeMreB3 wild type (cyan), SpeMreB3 D147E (pale blue), SpeMreB3 K174T (navy blue), SpeMreB3 S176D (light purple), and SpeMreB3 K174T/S176D (deep purple) in the presence of 2 mM Mg-ATP. (**H**) Time course plots of $P_i$ release of 3 μM SpeMreB5 wild type (red), SpeMreB3 T160A (orange), and SpeMreB5 D162S (pink) in the presence of 2 mM Mg-ATP.

**Figure S8. Sedimentation assay of the SpeMreBs.** Each SpeMreB was incubated after initiating polymerization and subjected to the ultracentrifugation of 436,000 × *g* for 120 min at 23°C. Precipitates were resuspended with water equivalent amount to the samples. For SpeMreB3, each fraction was diluted three times before the preparation of the sample for SDS-PAGE. Each fraction was loaded onto 12.5% Laemmli gel and stained with Coomassie brilliant blue to quantify the concentrations. (**A**) Sedimentation assay of 8 µM SpeMreB3 (left side of the lane M) and 3 µM SpeMreB5 (right side of the lane M) polymerized for 3 hours with the standard buffer in the presence ((+) ATP) or absence ((-) ATP) of Mg-ATP. Protein size standards are visualized in the lane M with the molecular masses of each band beside the band. (**B**) (left) Sedimentation assay of (top) 5 µM SpeMreB3 and (bottom) 1 µM SpeMreB5 (the minimum concentrations to determine each critical concentration) dependent on the polymerization time before the centrifugation. The SpeMreBs polymerized in buffer S for 1, 3, or 6 hours were subjected to the ultracentrifugation. (right) The pellet concentrations were quantified from the left gels and plotted over the incubation time as cyan and red crosses for SpeMreB3 and SpeMreB5, respectively. Error bars indicate SD from three repeated measurements. Symbols “n.s.” indicate *p*-values greater than 0.05 supported by Student’s *t*-test.

### Table S1

| Crystal | Nf-SpeMreB3 | SpeMreB3 AMPPNP |
|---|---|---|
| Space group | $P2_1$ | $P2_1$ |
| Cell dimensions | | |
| *a, b, c* (Å) | 52.4, 68.1, 54.6 | 50.3, 56.3, 120.5 |
| $\alpha, \beta, \gamma$ (deg) | 90.0, 91.7, 90.0 | 90.0, 90.6, 90.0 |
| Wavelength (Å) | 1.000 | 1.000 |
| Resolution (Å) | 54.5-1.90 (1.94-1.90) | 56.3-1.75 (1.78-1.75) |
| $R_{merge}$ | 0.094 (0.542) | 0.101 (0.570) |
| $CC_{1/2}$ | 0.996 (0.866) | 0.993 (0.658) |
| $I/\sigma I$ | 9.8 (2.7) | 6.8 (1.8) |
| Completeness (%) | 100.0 (100.0) | 97.2 (95.8) |
| Redundancy | 5.3 (5.2) | 3.3 (3.3) |
| Resolution range (Å) | 54.5-1.90 (1.95-1.90) | 51.0-1.75 (1.79-1.75) |
| No. of reflections working | 28,311 (2032) | 64,014 (4479) |
| No. of reflections test | 2001 (127) | 2006 (148) |
| $R_w$ (%) | 18.5 (22.1) | 17.4 (22.8) |
| $R_{free}$ (%) | 25.4 (28.1) | 21.4 (25.3) |
| Rms deviation bond length (Å) | 0.004 | 0.006 |
| Rms deviation bond angle (deg) | 0.614 | 0.891 |
| B-factors | | |
| Protein atoms | 27.4 | 25.1 |
| Ligand atoms | 20.4 | 17.9 |
| Solvent atoms | 28.5 | 26.0 |
| Ramachandran plot (%) | | |
| Most favored | 98.6 | 97.4 |
| Additionally allowed | 1.4 | 2.6 |
| Generously allowed | 0.0 | 0.0 |
| Disallowed | 0.0 | 0.0 |
| No. of protein atoms | 2519 | 5170 |
| No. of ligand atoms | 5 | 64 |
| No. of solvent atoms | 333 | 763 |

### Figure S1

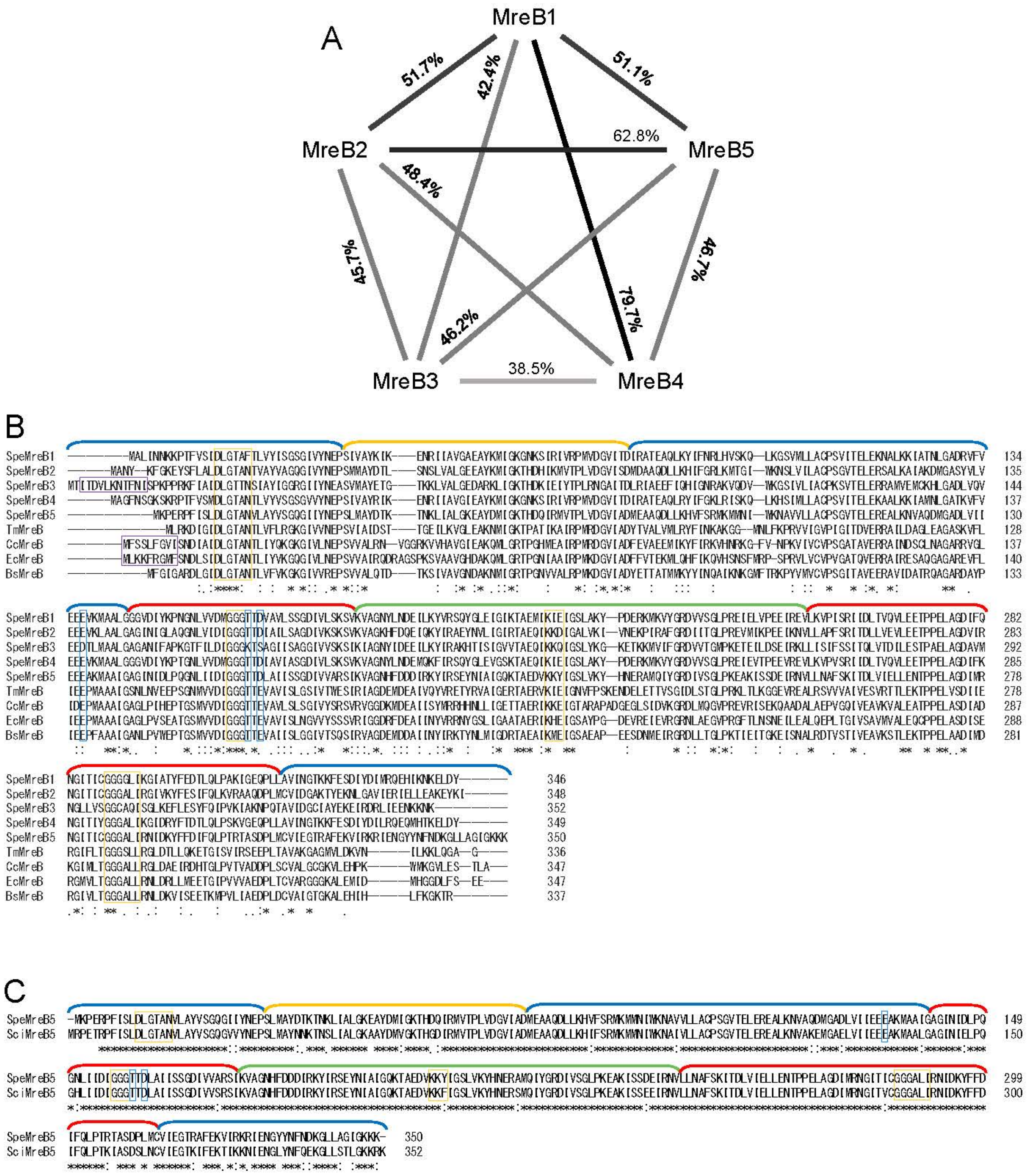
A
MreB1
MreB2
MreB3
MreB4
MreB5
51.7%
42.4%
51.1%
62.8%
48.4%
45.7%
46.7%
79.7%
46.2%
38.5%
B
SpeMreB1
SpeMreB2
SpeMreB3
SpeMreB4
SpeMreB5
TmMreB
CcMreB
EcMreB
BsMreB
134
135
144
137
130
128
137
140
133
282
283
292
285
278
278
287
288
281
346
348
352
349
350
336
347
347
337
C
SpeMreB5
SciMreB5
149
150
299
300
350
352

### Figure S2

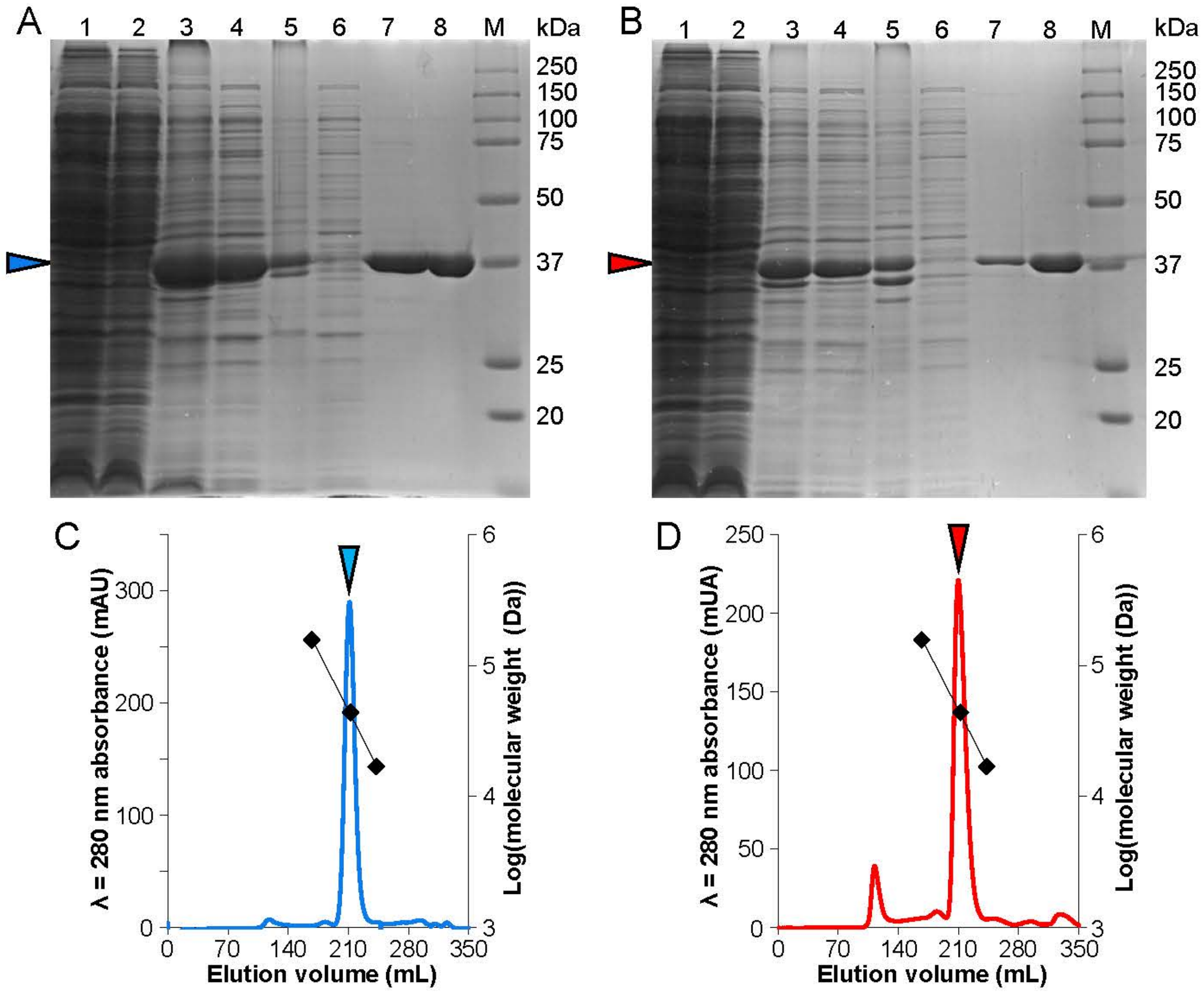
A
1 2 3 4 5 6 7 8 M kDa
250
150
100
75
50
37
25
20
B
1 2 3 4 5 6 7 8 M kDa
250
150
100
75
50
37
25
20
C
λ = 280 nm absorbance (mAU)
Log(molecular weight (Da))
0
100
200
300
3
4
5
6
0 70 140 210 280 350
Elution volume (mL)
D
λ = 280 nm absorbance (mUA)
Log(molecular weight (Da))
0
50
100
150
200
250
3
4
5
6
0 70 140 210 280 350
Elution volume (mL)

### Figure S3

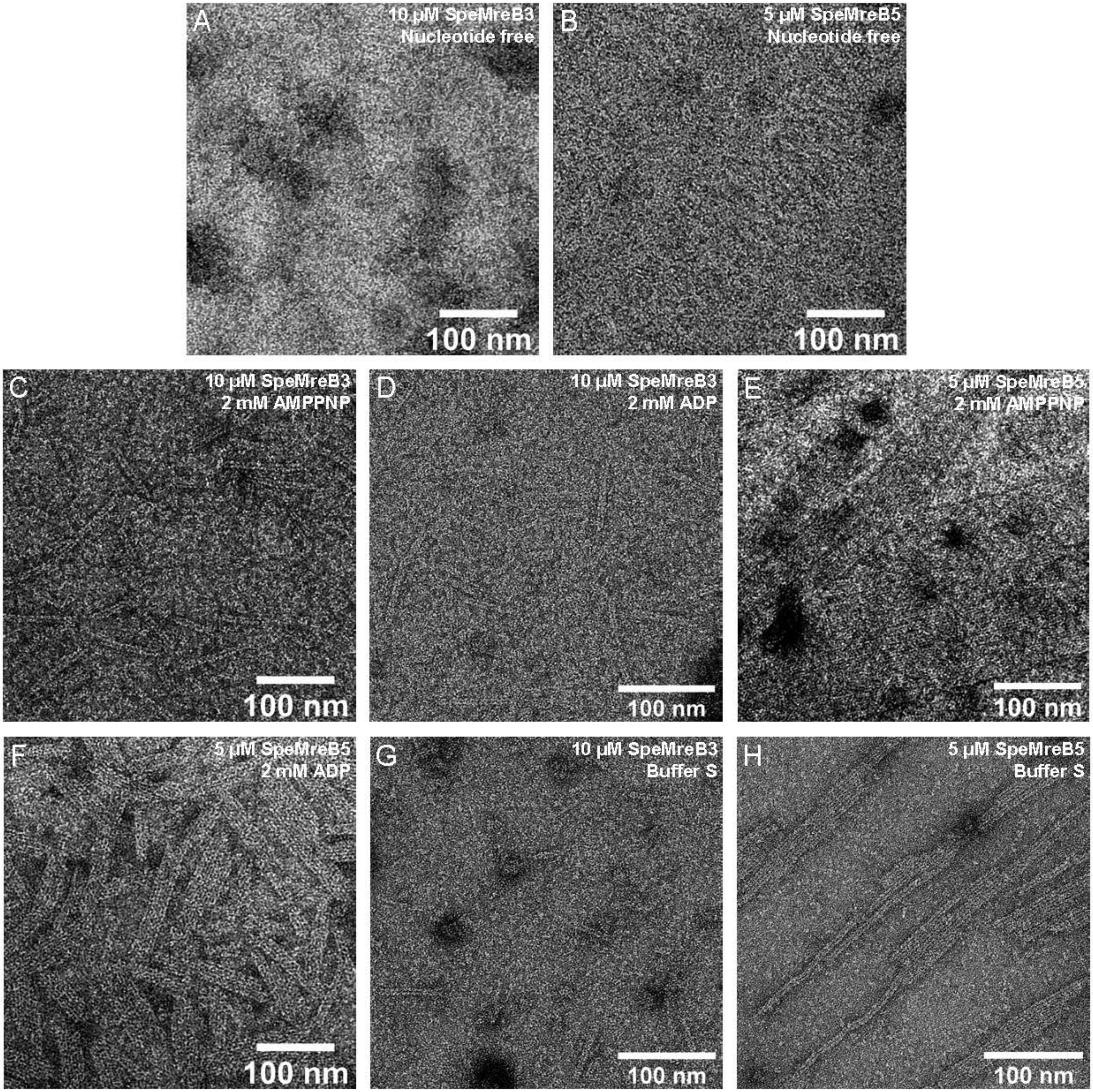
A
10 μM SpeMreB3
Nucleotide free
100 nm
B
5 μM SpeMreB5
Nucleotide free
100 nm
C
10 μM SpeMreB3
2 mM AMPPNP
100 nm
D
10 μM SpeMreB3
2 mM ADP
100 nm
E
5 μM SpeMreB5
2 mM AMPPNP
100 nm
F
5 μM SpeMreB5
2 mM ADP
100 nm
G
10 μM SpeMreB3
Buffer S
100 nm
H
5 μM SpeMreB5
Buffer S
100 nm

### Figure S4

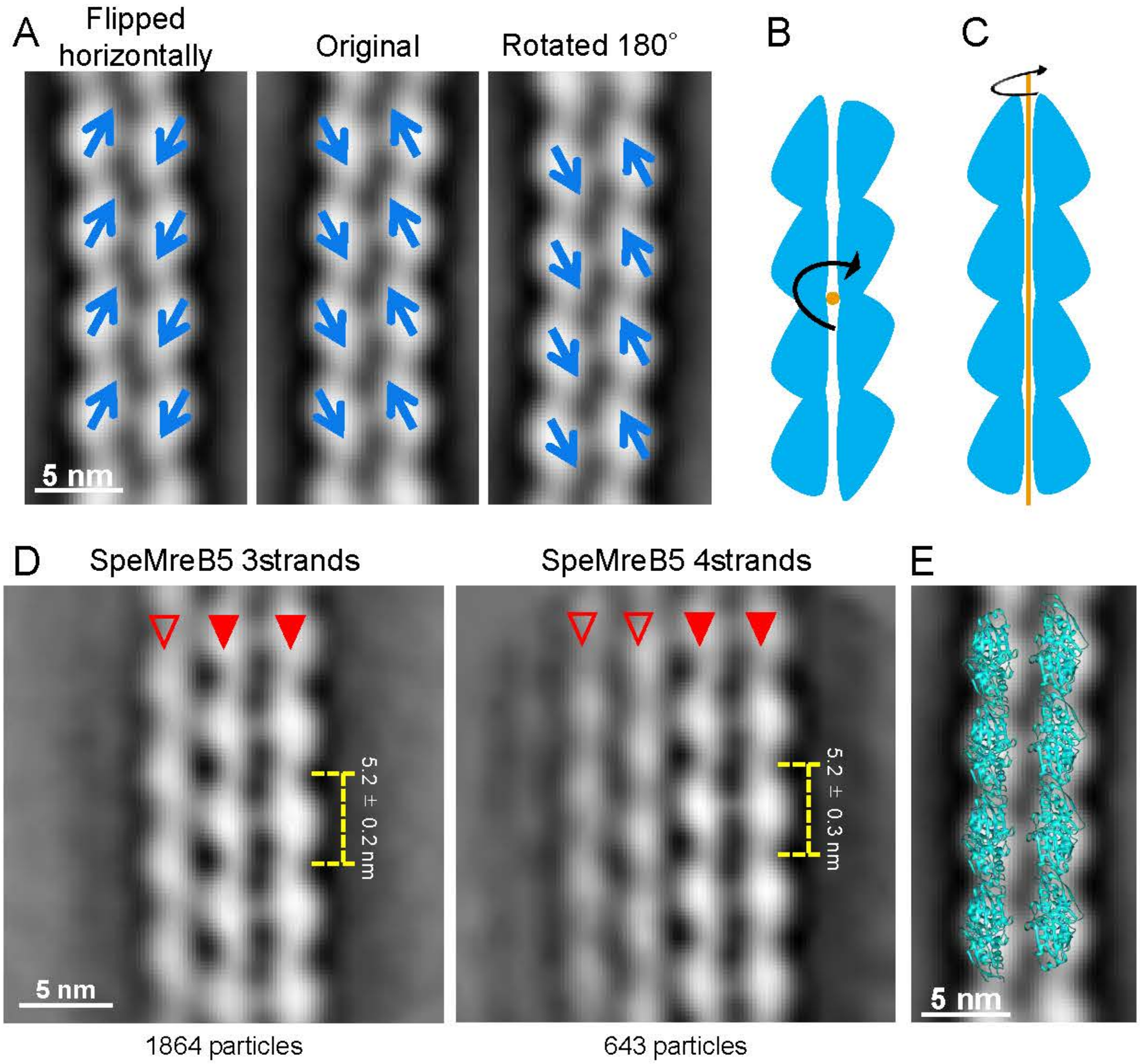

A
Flipped horizontally
Original
Rotated 180°
5 nm
B
C
D
SpeMreB5 3strands
SpeMreB5 4strands
5.2 ± 0.2 nm
5.2 ± 0.3 nm
5 nm
1864 particles
643 particles
E
5 nm

### Figure S5

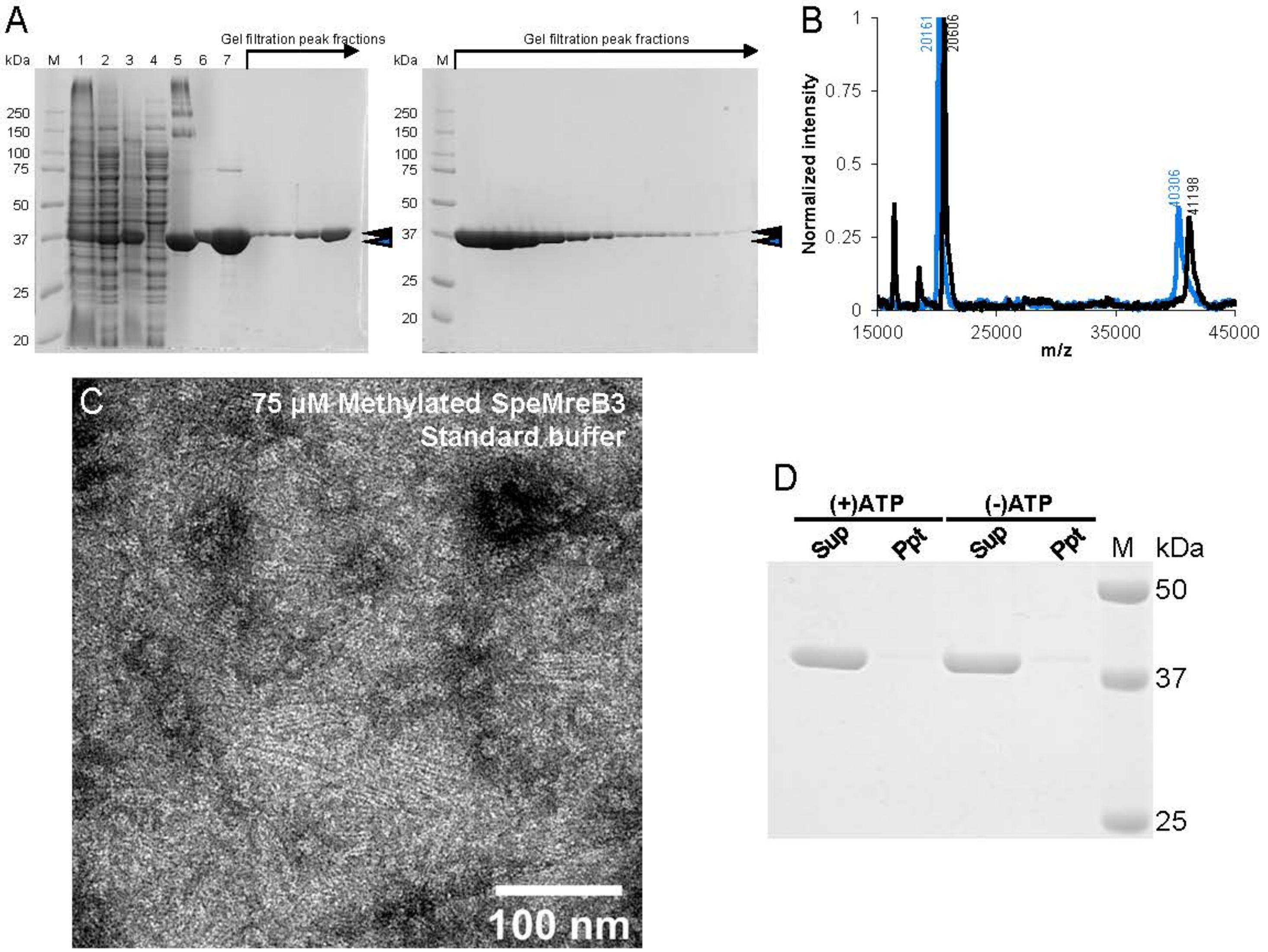

A
Gel filtration peak fractions
kDa M 1 2 3 4 5 6 7
250
150
100
75
50
37
25
20
Gel filtration peak fractions
kDa M
B
20161
20606
40306
41198
Normalized intensity
m/z
15000
25000
35000
45000
C
75 μM Methylated SpeMreB3
Standard buffer
100 nm
D
(+)ATP
(-)ATP
Sup
Ppt
Sup
Ppt
M
kDa
50
37
25

### Figure S6

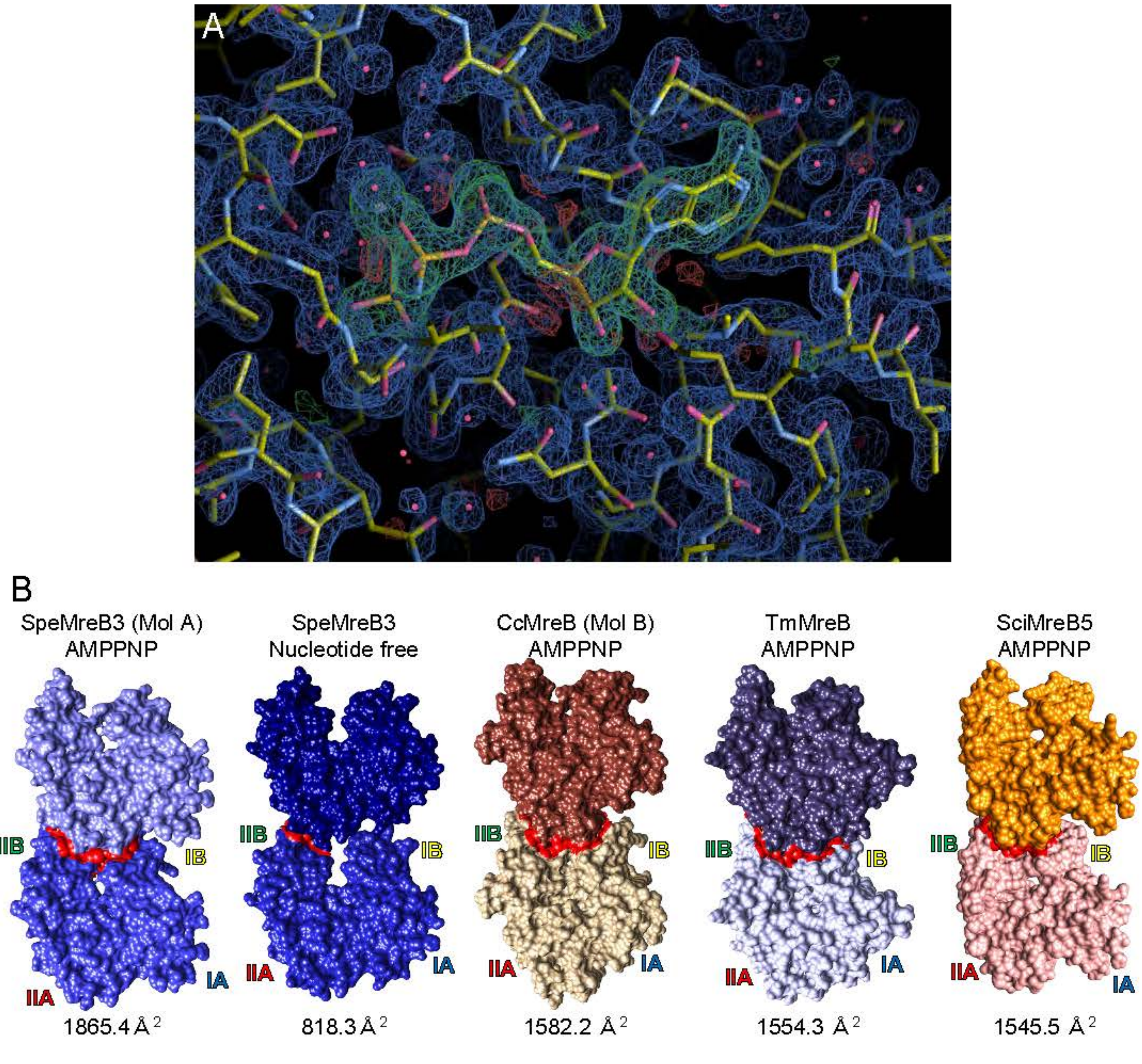

A
B
SpeMreB3 (Mol A)
AMPPNP
SpeMreB3
Nucleotide free
CcMreB (Mol B)
AMPPNP
TmMreB
AMPPNP
SciMreB5
AMPPNP
IIB
IB
IIA
IA
1865.4 Å 2
818.3 Å 2
1582.2 Å 2
1554.3 Å 2
1545.5 Å 2

### Figure S7

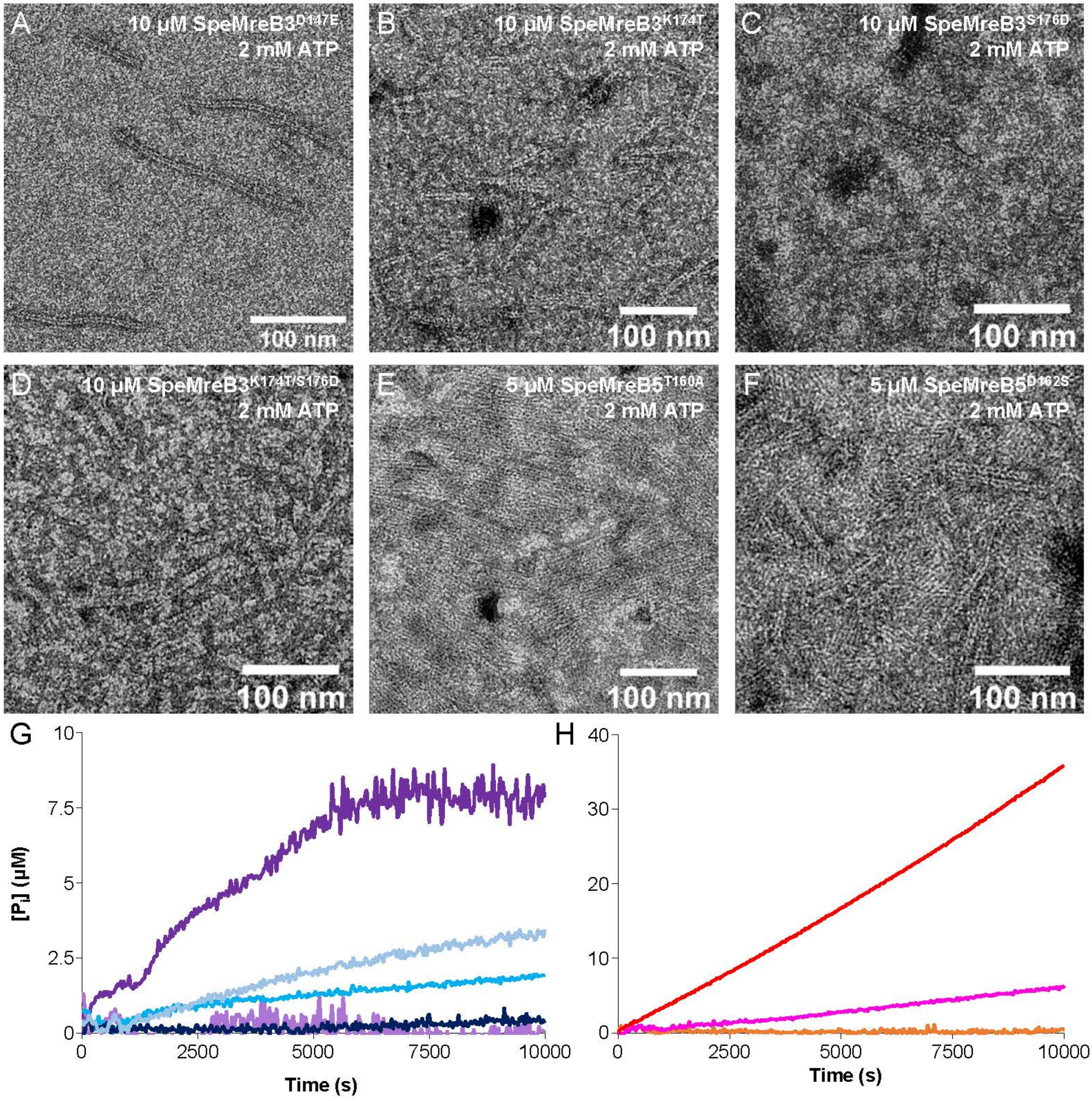

A
10 µM SpeMreB3D147E
2 mM ATP
100 nm
B
10 µM SpeMreB3K174T
2 mM ATP
100 nm
C
10 µM SpeMreB3S176D
2 mM ATP
100 nm
D
10 µM SpeMreB3K174T/S176D
2 mM ATP
100 nm
E
5 µM SpeMreB5T160A
2 mM ATP
100 nm
F
5 µM SpeMreB5D162S
2 mM ATP
100 nm
G
[Pi] (µM)
0
2.5
5
7.5
10
0
2500
5000
7500
10000
Time (s)
H
0
10
20
30
40
0
2500
5000
7500
10000
Time (s)

### Figure S8

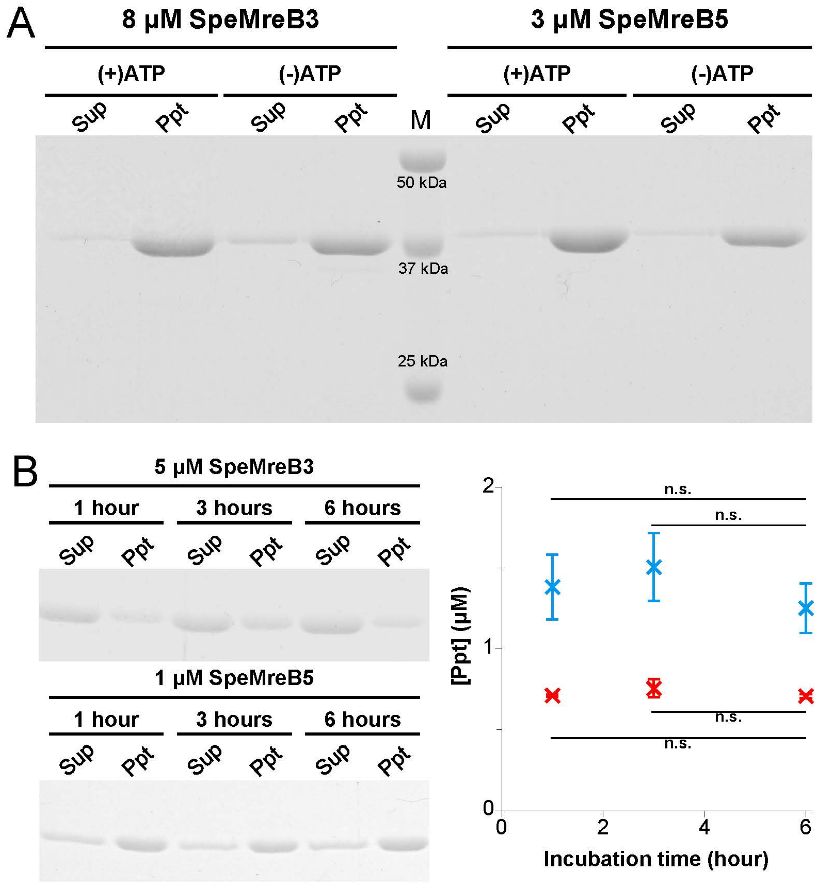
A
8 µM SpeMreB3
3 µM SpeMreB5
(+)ATP
(-)ATP
(+)ATP
(-)ATP
Sup
Ppt
Sup
Ppt
M
Sup
Ppt
Sup
Ppt
50 kDa
37 kDa
25 kDa
B
5 µM SpeMreB3
1 hour
3 hours
6 hours
Sup
Ppt
Sup
Ppt
Sup
Ppt
1 µM SpeMreB5
1 hour
3 hours
6 hours
Sup
Ppt
Sup
Ppt
Sup
Ppt
n.s.
n.s.
n.s.
n.s.
[Ppt] (µM)
0
1
2
0
2
4
6
Incubation time (hour)
